# Natural selection driven by DNA binding proteins shapes genome-wide motif statistics

**DOI:** 10.1101/041145

**Authors:** Long Qian, Edo Kussell

**Affiliations:** Department of Biology and Center for Genomics and Systems Biology, New York University, 12 Waverly Place, New York, New York, 10003, USA; Department of Physics, New York University, 4 Washington Place, New York, 10003, USA

## Abstract

Ectopic DNA binding by transcription factors and other DNA binding proteins can be detrimental to cellular functions and ultimately to organismal fitness. The frequency of protein-DNA binding at non-functional sites depends on the global composition of a genome with respect to all possible short motifs, or *k*-mer words. To determine whether weak yet ubiquitous protein-DNA interactions could exert significant evolutionary pressures on genomes, we correlate *in vitro* measurements of binding strengths on all 8-mer words from a large collection of transcription factors, in several different species, against their relative genomic frequencies. Our analysis reveals a clear signal of purifying selection to reduce the large number of weak binding sites genome-wide. This evolutionary process, which we call *global selection*, has a detectable hallmark in that similar words experience similar evolutionary pressure, a consequence of the biophysics of protein-DNA binding. By analyzing a large collection of genomes, we show that global selection exists in all domains of life, and operates through tiny selective steps, maintaining genomic binding landscapes over long evolutionary timescales.

## 1 Introduction

DNA sequences encode information that is read and interpreted through molecular binding by proteins including transcription factors (TFs), nucleosomes, the RNA polymerase complex, and DNA replication machinery. A DNA binding factor must discriminate a small number of target sites from the set of all possible loci genome-wide. While some DNA binding proteins exhibit exquisite specificity, the majority display fuzziness in their binding preferences [1, 2], and DNA binding often relies on cooperative interactions and chromatin accessibility to increase specificity [3]. Since accessible genomic regions constitute a large amount of DNA (~450 Mb in the human genome) and a substantial proportion of transcribed regions [4], ectopic DNA binding can interfere with multiple processes, including transcription, replication, and nucleosome positioning. Moreover, it titrates copies of DNA binding proteins away from functional sites, reducing the efficiency of gene regulation. Since for any given TF there are exponentially more weak binding sites than strong ones [1, 2], weak ectopic binding to a large number of sites genome-wide is potentially more detrimental to cellular functions than strong ectopic binding which may be comparatively rare. It should therefore be beneficial for genome sequences to evolve to reduce the frequency of non-functional binding sites genome-wide.

Due to the large number of loci involved, such *global selection* on genomes is expected to involve tiny selective coefficients that may be difficult to detect by traditional methods. Indeed, previous studies have identified only a handful of binding motifs that appear to be globally selected against, mainly in bacteria, including promoter elements [5, 6], transcription/translation boundary signals [7, 8], and restriction sites [9, 10]. Other aspects of genome-wide composition have been extensively studied, including global G/C content and codon usage [11, 12, 13, 14, 15, 16]. Seminal work by Karlin and co-workers indicated that dinucleotide frequencies differ between species [17, 18], and more recently, further differences at the level of longer *k*-mer words have been detected [19, 20]. While these compositional differences could modulate genome-wide binding, there is little consensus on whether mutational biases, drift, or natural selection are their major driving force [11, 12, 21, 22, 23, 14, 24, 25]. It remains largely unknown whether genome sequences have been substantially shaped by DNA binding-related evolutionary pressures.

Here, we demonstrate that the distinct set of DNA binding proteins coded in each species’ genome imposes a large set of global, evolutionary pressures that shape genome-wide motif composition. By correlating *in vitro* measurements of DNA binding with genome-wide word statistics, we show that genomes have evolved to reduce the occurrence of weak binding motifs. We introduce an evolutionary model of global selection, and use it to infer selective coefficients and to deduce the evolutionary timescales of global adaptation across all domains of life.

## 2 Results

### 2.1 Genomic binding landscapes of transcription factors

To investigate the impact of DNA binding factors on genomic binding landscapes, we correlated *in vitro* datasets on protein-DNA binding specificities against genomic sequence composition. We used the UniPROBE datasets [26], which are based on a protein-binding microarray that measures the binding of a protein to every possible 8-mer sequence (total 32, 896). We initially studied the mouse dataset [1] in which binding of 109 TFs to each 8-mer on the microarray was determined. For each TF, the log binding intensity values were centered to the median, and normalized by their dispersion, yielding a binding score *b*_*i*_ for every possible 8-mer word *i*. Fig. 1A shows the distribution of binding scores for a single TF (Mafk) across all 8-mers (see File S1 for all TFs). Words in the positive or negative tails correspond to very strong or very weak binding, respectively. For most TFs, the majority of words lie along a continuum of binding levels without substantial gaps, consistent with previous observations that TFs typically exhibit degeneracy of their target preferences [27, 28].

**Figure 1:**
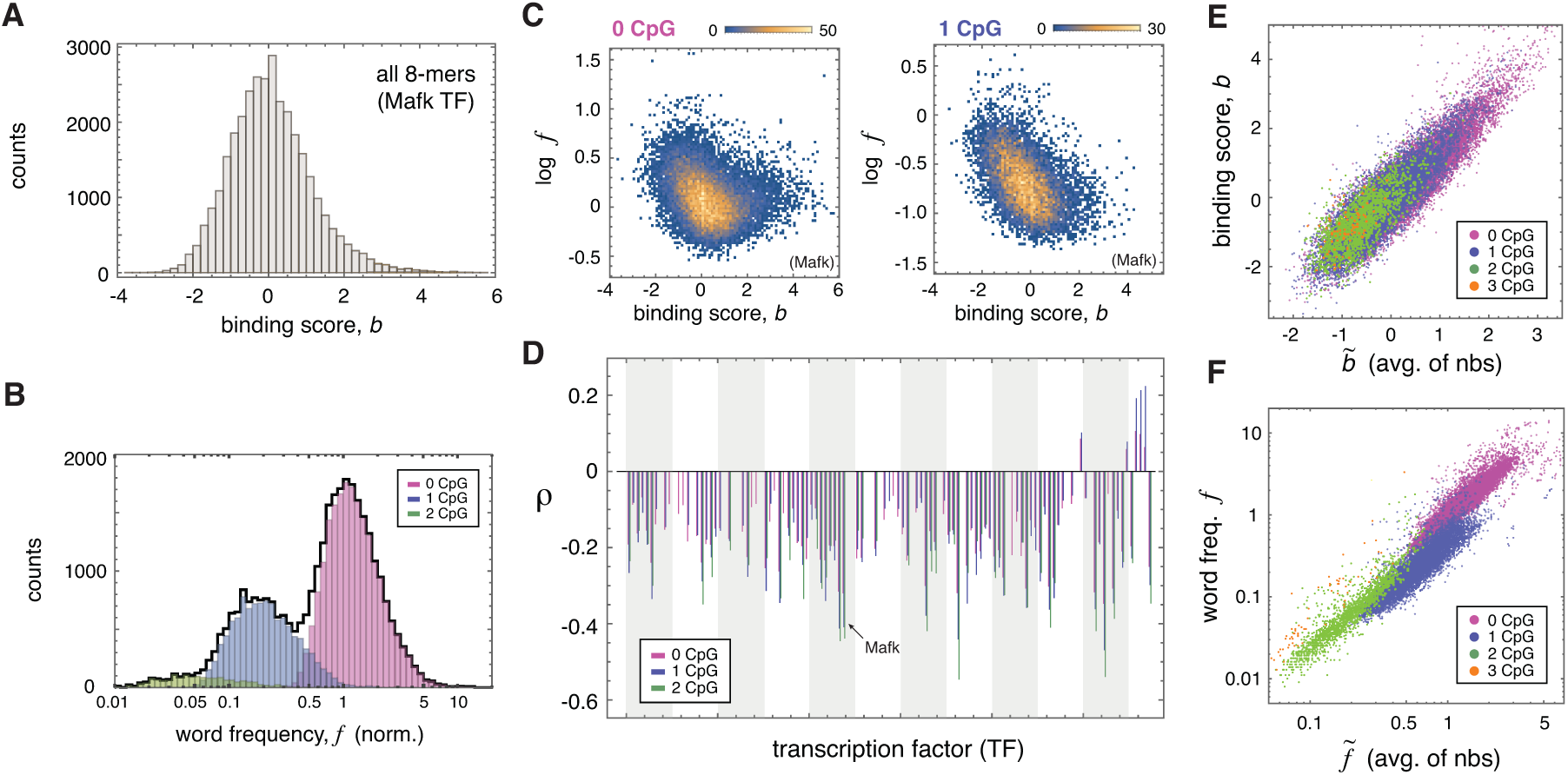
Mouse genomic binding landscape and *in vitro* DNA binding measurements. **A**. Distribution of binding scores *b*_*i*_ for the Mafk TF over all 8-mer words *i*. **B**. Distribution of 8-mer word frequencies *f*_*i*_ in mouse introns (black curve); *f*_*i*_ are shown normalized with respect to expectation based on genome-wide nucleotide composition. Words are separately histogramed according to their CpG counts (colored bars). **C**. Correlation of *b*_*i*_ and log *f*_*i*_ for the Mafk TF in separate CpG categories. Color indicates density of points. **D**. Correlation coefficients (Spearman’s *ρ)* of *b*_*i*_ vs. *f*_*i*_ are shown for each mouse TF, using all wea*k*-binding words *(b_i_ <* 0) separately computed conditioned on word CpG content. Bars are shown only for statistically significant correlations, with p-value < 10^−6^. **E**. Binding scores of words *(b_i_)* are correlated with the average binding score of their mutational neighborhoods 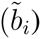; results shown for Mafk, 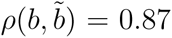. **F**. Correlation of *f*_*i*_ and 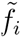 over all 8-mer words *i* for mouse introns; words are colored according to their CpG content, and frequencies are normalized as in **B & C**.

We analyzed the intron regions of the mouse genome, which constitute ~750Mb or nearly 30% of the total genomic DNA. Introns are ideal for detecting binding-related pressures because (i) they are largely devoid of locus-specific selective pressures (as found e.g. in exons) which confound detection of global effects, and (ii) they reside in genic regions and are thus generally accessible to binding factors. Simply correlating *in vitro* binding scores with genomic word frequencies, however, is highly misleading due to two effects. First, the nucleotide composition (G/C content) of the genome is a major predictor of word usage, and may be influenced by mutational biases, drift, and selection [22, 24]. TFs that bind words composed of more (less) frequent nucleotides are likely to exhibit positive (negative) correlations between binding scores and genomic *k*-mer frequencies regardless of evolutionary history. Thus, even if G/C content itself were evolutionarily shaped by binding-related global pressures, correlations between raw word frequencies and TF binding scores cannot be used as evidence. In our analysis, we therefore used relative word frequencies, i.e. normalized by expectation based on genome nucleotide composition. Second, the distribution of *k*-mer frequencies in mouse introns is bimodal due to differences in words’ CpG content (Fig. 1B). In vertebrates and plants, the dinucleotide CpG is hypermutable (e.g. in mice its mutation rate is ~10 times the average point mutation rate) causing genome-wide depletion of words that contain it [29]. To account for this large effect, which masks smaller differences among words, we correlated *k*-mer binding scores vs. relative word frequencies separately for words with different CpG content.

The example in Fig. 1C shows pronounced negative correlations in each CpG category, indicating that the stronger binding a word, the less frequently it is used in the genome relative to expectation. For most TFs, we observed that words with below-average binding (b < 0) exhibit highly significant negative correlations (Fig. 1D). For words with above-average binding (b > 0), both positive and negative correlations were found, depending on the TF (Fig. S1). Correlating binding scores of all words against their relative frequencies yielded negative cor-relations in each CpG category for a majority of TFs (Fig. S1). Similar results were obtained in worm (21 TFs), fly (14 TFs), human (8 TFs), and yeast (89 TFs) genomes (Fig. S2, Table S1, and File S1). We conclude that statistically significant correlations exist between binding scores and genomic relative word frequencies, and that in general, weak binding words are avoided compared to even weaker binding words.

### 2.2 A genomic hallmark of global selection

We sought a more general method that could be applied in the absence of *in vitro* measurements to detect global selection due to DNA binding-related pressures. We noticed that, consistent with the biophysics of protein-DNA interaction [30], TF binding scores of words that differ at a single nucleotide are strongly correlated: for each word *i* we plot its binding score *b*_*i*_ vs. the average binding score 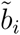 of its ‘mutational neighbors’ - all words that differ from *i* at a single nucleotide (Fig. 1E & File S1). For all 241 TFs that we analyzed, we found a general statistical rule that *similar words have similar binding strengths*. Therefore, if genomes have adapted globally under DNA binding pressures, then we should detect a strong correlation between the frequencies of similar words, since *similar words would be under similar pressures*. In Fig. 1F we show the result for the mouse genome, where for each 8-mer word *i*, its frequency *f*_*i*_ is plotted vs. the average frequency 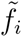 of its mutational neighbors. Consistent with the hypothesis, we observed a strong correlation of *f*_*i*_ and 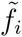 (*ρ >* 0.85 within each CpG group).

We tested for word-neighbor correlations in a large collection of fully sequenced genomes spanning all domains of life. Word frequencies were measured separately in exon and intron regions, and normalized by using appropriate null models that account for context-dependent mutational biases and other compositional effects. For exons, we used synonymous codon and dicodon shuffling schemes to construct partially randomized DNA sequences that preserves amino acid sequences, genomic codon biases, and nucleotide base composition. The dicodon shuffling scheme additionally preserves the frequencies of all *k*-mers for *k* ≤ 4. The randomized sequences were scanned to determine expected word frequencies for exons. For introns, we computed expected word frequencies based on genome-wide nucleotide (1-mer), dinucleotide (2-mer), or trinucleotide (3-mer) frequencies.

All tested genomes (947 bacterial/archaeal genomes, 1304 eukaryotic chromosomes from 75 species) exhibited striking correlation between *f*_*i*_ and 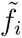 in exons (Figs. 2A, S3, S4, S7, S23) and in introns (Figs. S5, S6A, S6B) for each of the null models. Normalized word frequencies correlated strongly between exon and intron regions (Fig. S8). We separately analyzed all DNaseI hyper-sensitive regions of the human genome, which are verified binding-accessible regions, and these exhibited similar word-neighbor correlations (Fig. S22). Eukaryotic genomes were further analyzed on a chromosome-by-chromosome basis. In human chromosomes, for example, the overall shape of the *f* vs. 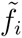 plots from exons is qualitatively similar across chromosomes (Fig. 2B & Fig. S9). Comparing relative word frequencies *f*_*i*_ between different chromosomes, we found high correlation coefficients (ρ > 0.9) for most chromosome pairs within each genome (Table S4). Deviations were observed for short or Y chromosomes (Fig. 2B & Fig. S10A), due to insufficient word sampling as well as strong genetic linkage, discussed below.

**Figure 2:**
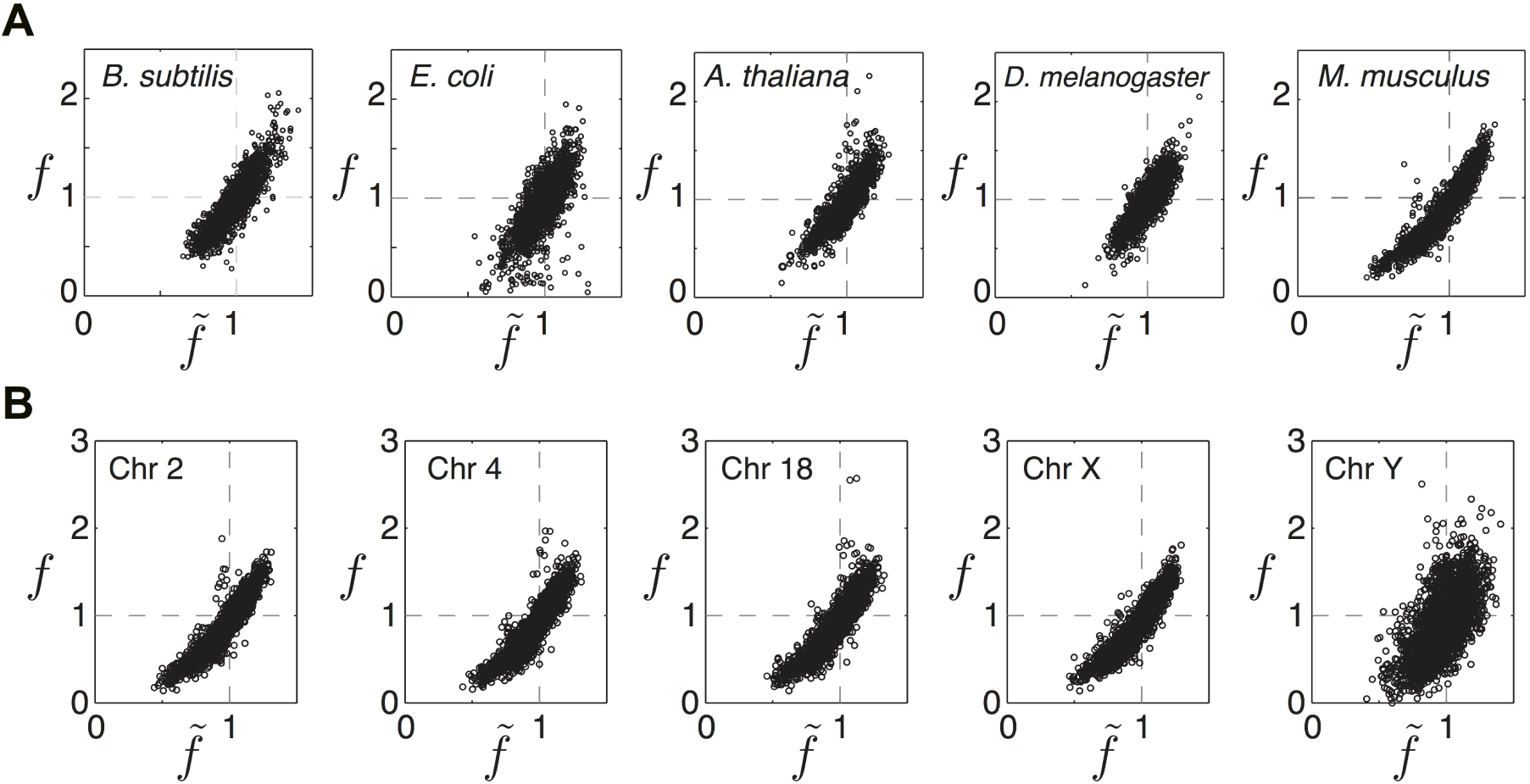
Word-neighbor correlations are a hallmark of global selection. **A**. Word frequency plots of *f*_*i*_ vs. 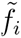 in exons of representative bacterial and eukaryotic genomes. Points correspond to all possible 6 bp words. Frequencies are shown relative to the null expectation from synonymous codon-shuffling, where a value of 1 means the observed frequency is equal to the expected frequency. For eukaryotes, frequencies were computed across all chromosomes. **B**. Word frequency plots from exons of individual human chromosomes (see Fig. S9 for all data).

### 2.3 Mathematical model of global selection

We examined the consequences of a global selective process acting differentially on words across a genome. We represented a genome by its *k*-mer frequency vector *f*_*i*_, where each word *i* experiences evolutionary pressure according to a selective coefficient *s*_*i*_. A population of genomes evolves by successive rounds of mutation and selection, where mutations cause random changes to the word vectors with rate *u* (per bp), and genomes reproduce proportionally to the total fitness of their words. The model admits a unique equilibrium solution in the large population limit, which expresses the stable word frequencies in terms of the selective coefficients and mutation rate *(Supplementary Text)*. Conveniently, it is possible to invert this relation, and for small *u* we obtain

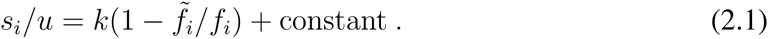

Selective coefficients relative to *u* are determined up to an additive constant by the ratio of a word’s frequency and the average frequency of its mutational neighbors. Moreover, this relation generalizes to include biased mutation rates *(Supplementary Text)*. Fig. 3A-C shows examples of the model solution when selective coefficients are randomly sampled from a normal distribution with standard deviation *σ*. Words experiencing similar pressures fall on the lines in the 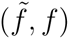-plane defined by (1) (Fig. 3A). This result encapsulates a basic insight of our analysis: the genome-wide frequency of a word says little if anything about the global pressure it experiences. Words can be under-or over-represented simply because their mutational neighbors are under negative or positive pressures, respectively. Indeed, selective coefficients can only be inferred when a word is viewed relative to its mutational neighbors.

**Figure 3:**
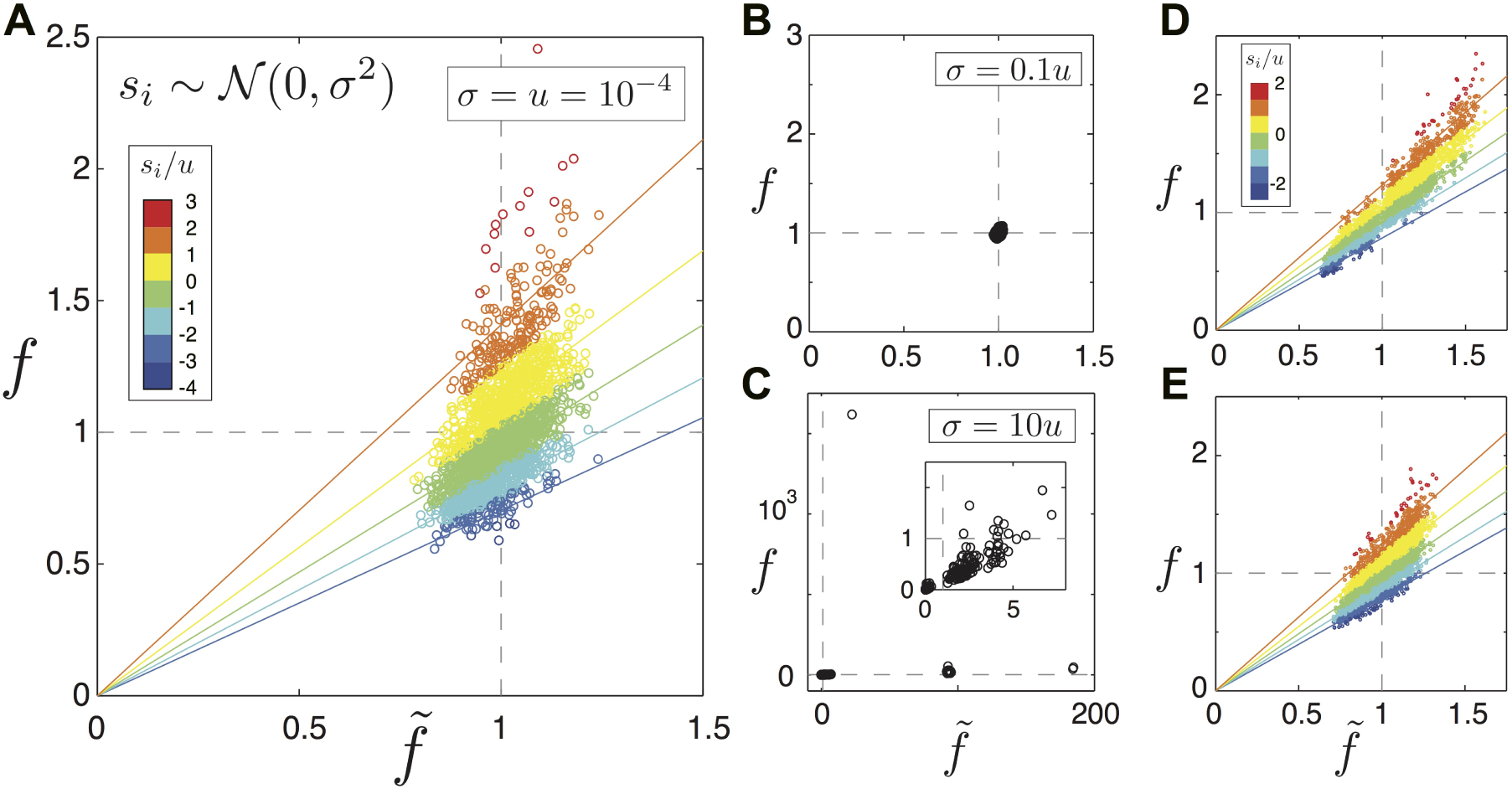
Mathematical model of global selection. **A-C**. Solution of the model at mutation-selection balance. Plots in the 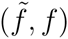-plane of the equilibrium word frequencies for independent, normally-distributed selective coefficients *s*_*i*_ ~ 𝒩(0, σ^2^), using the indicated values of *σ.* We numerically solved the model’s eigenvector equation *(Supplementary Text*, Eq. 2.2) to determine the equilibrium word frequencies. Color shows different bins of selective coefficients, and predicted equal-pressure lines are drawn using each interval’s average pressure. Inset in C shows a zoomed view. Parameters were *k* = 6 and *u* = 10^−4^, and frequencies are shown relative to the neutral expectation *4*^*-k*^. **D**. Solution when using strongly correlated pressures on neighboring words. **E**. Correlation is reduced by shuffling 50% of words’ pressures from D.

When selection is weak relative to mutation (σ *< u*, Fig. 3B) the word cloud collapses toward a point in which all words are effectively neutral, while under strong selection (σ *> u*, Fig. 3C) the word with the maximum selective coefficient dominates the distribution. Only in the intermediate regime *(σ* ≃ *u*, Fig. 3A) does the solution take the form of an extended word cloud, with frequencies varying from approximately twofold avoidance to twofold enrichment. A pronounced positive correlation between *f*_*i*_and 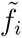 is seen, despite the fact that all words experience independent random pressures. This correlation results from selection, which modulates the frequency of neighbor words in order to alter mutational fluxes into words under selection (similar results are obtained with a skewed pressure distribution, Fig. S11). The correlation of frequencies can be increased further by introducing positive correlations in the pressures on similar words, resulting in an extended, rotated word cloud (Fig. 3D,E), while the equal-pressure lines remain unchanged.

### 2.4 Distribution of global selective coefficients in genomes

We applied the mathematical model to infer the global selective coefficients in each genome. Word frequencies *f*_*i*_ were counted separately in introns and exons, and expected frequencies 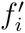 were obtained using the null models described above. We measured global selective coefficients using the *excess pressures* 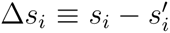, where *s*_*i*_ and 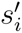 were separately inferred from *f*_*i*_ and 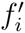, respectively, using Eq. 1. By using excess pressures *∆s*_*i*_ rather than *s*_*i*_, we measure the strength of selection needed to shift word frequencies to their observed values from their expected values based on the null models *(Supplementary Text)*. For example, mutational biases such as CpG hypermutation and other less pronounced biases influence the distribution of dinucleotides observed in the genome. By measuring excess pressures relative to the 2-mer null model, we measure only the additional selective pressures that are not already accounted for by dinucleotide composition. In Fig. 4 the distribution of selective coefficients is shown for several genomes (see Figs. S3-S7 for additional genomes and null models). In all cases, the bulk of the distribution has a width comparable to *u*. The selective coefficients *∆s*_*i*_ measured on different chromosomes were strongly correlated, with overall very similar distributions (Fig. S9). Importantly, Eq. 1 allows us to determine the pressures on each word in spite of pressures acting on their mutational neighbors. Consistent with our primary hypothesis, we find that the selective pressures on similar words are indeed highly correlated (Fig. S13).

**Figure 4:**
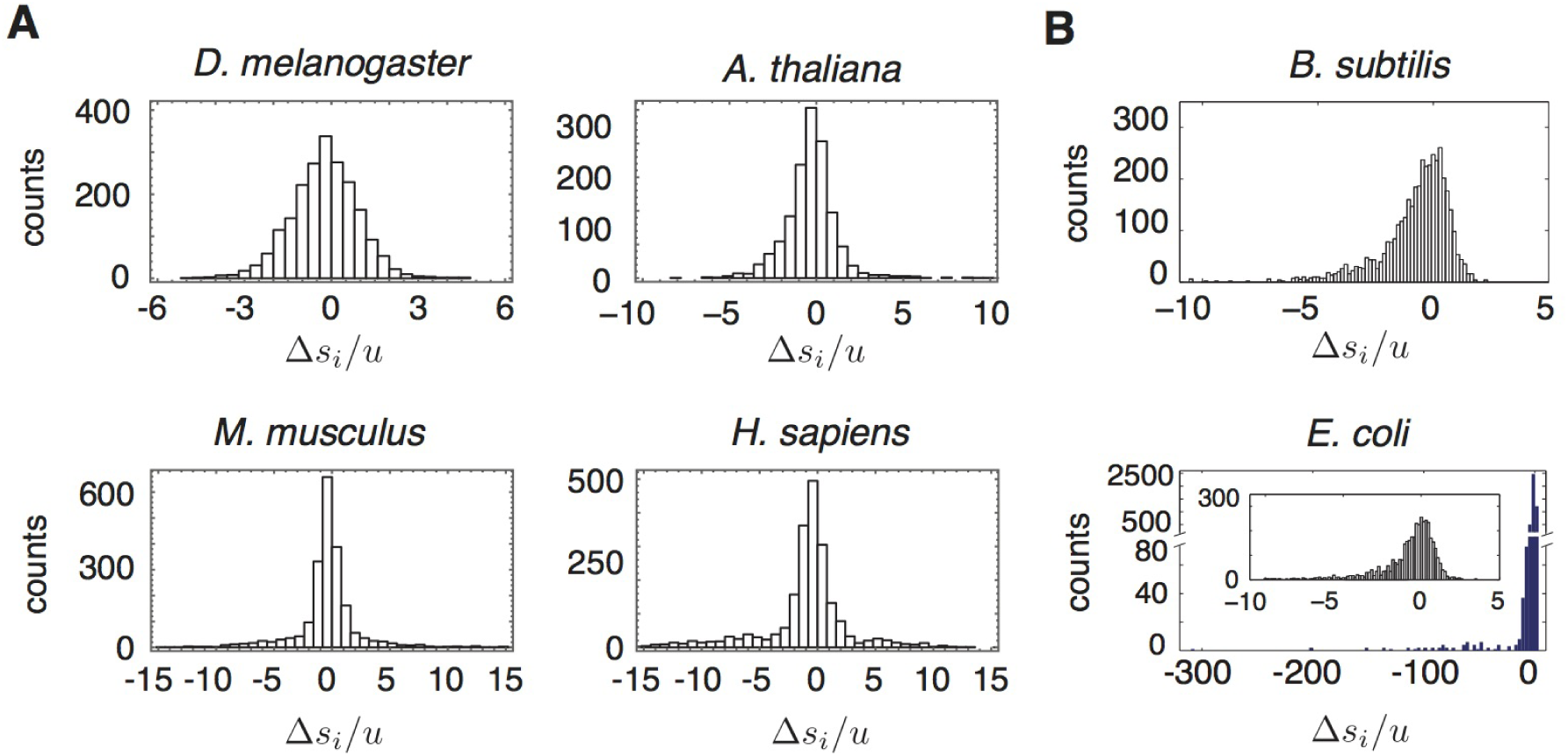
Distribution of global selective coefficients in different genomes. **A**. Selective coefficients of 6-mers in four eukaryotic species. Intron data was used and ∆*s*_*i*_ values are given with respect to the 2-mer null model. **B**. Selective coefficients of 6-mers measured in two bacterial species. Exon data was used and ∆*s*_*i*_ values are given with respect to the synonymous codon-shuffling model. In *E.coli* an inset shows the bulk of the distribution, since a small number of words including known restriction sites have large negative coefficients.

### 2.5 Neutral mechanisms are unable to account for observed word frequency distributions and word-neighbor correlations

To determine whether non-selective mechanisms, such as mutational biases and repeat expansion, can account for the observations we ran a wide range of tests. Controls for mutational biases included performing an analysis of variance on *k*-mer frequencies using their dinucleotide and trinucleotide composition (Table S5), analysis of word-neighbor correlations using regression residuals (Fig. S16), a dicodon shuffling scheme that accounts for mutational biases in bacteria (Fig. S23), and explicit incorporation of mutational biases into evolutionary models (Fig. S12). In each of these tests, mutational biases were unable to account for the observed word frequency distributions and word-neighbor correlations (see *Supplementary Text* for a full discussion). Since large eukaryotic genomes have a substantial amount of repeat-derived sequence, we tested whether word-neighbor correlations might arise from a balance between amplification of specific classes of mobile and/or repeat-containing elements and mutational degradation. We analyzed the repeat-masked sequence of the human genome, a procedure that removes approximately 45% of the sequence (Fig. S21), and found that it exhibited strong word-neighbor correlations (*r* ≥ 0.92 for all null models, Fig. S21C). Repeat expansion is therefore not responsible for word-neighbor correlations, and word frequencies were strongly correlated between repeat-masked and repeat-derived regions (Fig. S21B). A different possibility is that due to ubiquitous small insertions and deletions (indels) occurring in all genomic regions, word composition could be determined by slippage-based mutational mechanisms, a phenomenon known as ‘cryptic simplicity’ [31]. While repeat-masking cannot detect this finer-scale process, it is well-known from comparative genomics and mutation accumulation studies in different species that indels occur between 0.03 - 0.13 times as frequently as point mutations [32, 22, 33, 34, 35]. The mathematical model shows that processes that change word frequencies at much slower rates than the point mutation rate cannot yield the observed word-neighbor clouds (Fig. 3B). These results indicate that neither mutational biases nor neutral processes involving repetitive DNA can account for the ubiquitous word-neighbor correlations observed.

### 2.6 Ancient phylogenetic signal of global selection

Evidence of a persistent global evolutionary process acting on *k*-mers can be found in a principal component analysis (PCA) we performed on intronic relative word frequencies (2-mer null) across eukaryotic species (Fig. 5). Chromosomes within a single genome form tight clusters that demarcate species. Although major groups of eukaryotes can be clustered to a certain extent using genome-wide di-, tri-, and tetra-nucleotide frequencies [17, 19] as well as codon biases [12], we achieved a significantly higher resolution using data exclusively from the intron regions of individual chromosomes. Closely related species were spatially proximate, with plants, invertebrates, and vertebrates following different directions in the PCA space (Fig. S17). As discussed below, these remarkable phylogenetic signals are not explained by neutral divergence processes (see Discussion).

**Figure 5:**
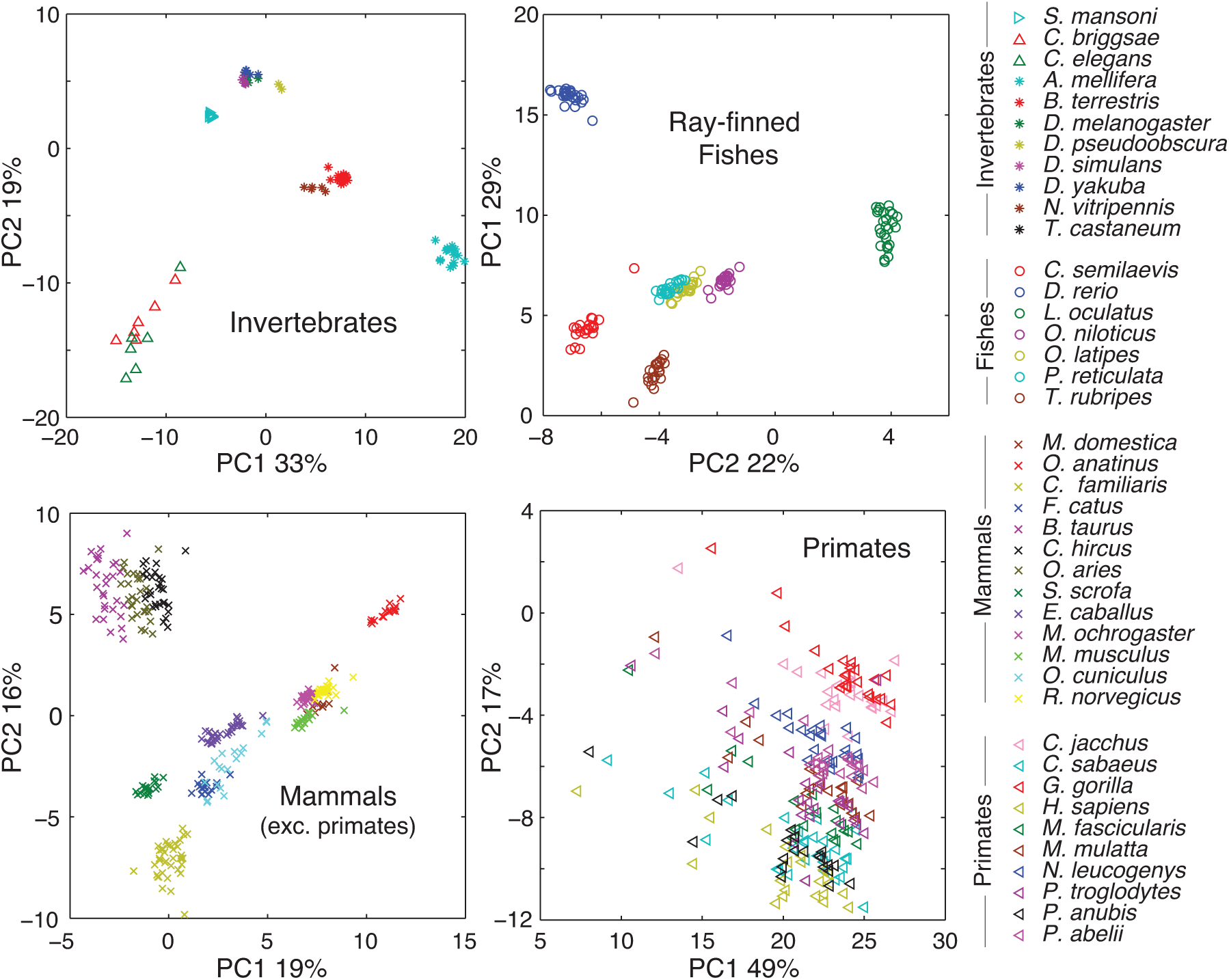
Cross-species comparison of intron word frequencies using principle components analysis (PCA). Each point in the PCA plots corresponds to a single chromosome projected on the two principal axes. Principal axes are computed using the 6-mer relative word frequency vectors (2-mer null model) of all chromosomes shown in each plot. See Table S3 for further species information.

Our analysis indicates that primates experienced a ‘jump’ in global pressures from the rest of the mammals (Fig. S18); while within this group, no species clusters were detected (Fig. 5), indicating that the global pressures have not significantly changed since the last common ancestor. In prokaryotes, phylogenetic signals are generally very weak or non-existent when assessed across all genomes (Fig. S19), but may persist at the genus and species level (Fig. S20), indicating that over the longest evolutionary timescales global pressures can change so extensively that the most ancient parts of the signal are extremely faint and difficult to detect.

### 2.7 Evolutionary dynamics of global selection

Our mathematical model is only valid under mutation-selection balance, hence the selective coefficients that we infer (Fig. 4) correspond to the magnitude of purifying selection necessary to maintain genome-wide motif statistics over evolutionary timescales. Since the effect sizes on individual words are tiny, we asked whether the global fitness differences between individuals are sufficiently large to constitute non-negligible selective differences, i.e. larger than ~ *1/N*_*eff*_. If two individuals differ at l sites, or *kl* words, and each word contributes an average effect size 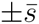, their global fitness difference 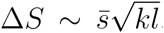. In human populations, pairs of individuals differ at 4.6 x 10^6^ sites on average [36], and taking *k* = 6 and 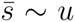 (Fig. 4), with *u* = 10^−8^ [37], we have *∆S* ≈ 5 x 10^−5^. Effective global selection thus requires *N*_*eff*_ ≳ 10^4^. In *E. coli*, isolates differ on average at 1 - 2% of sites [38], or approximately 7 x 10^4^ sites. Taking 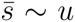, and using *u* = 2.2 x 10-^10^ [33], we find *∆S* ≈ 1.4 x 10-^7^, hence global selection requires *N_eff_ ≳* 10^7^. The per-word selective coefficients we measured in these genomes are thus sufficient to maintain genome-wide statistics given the present levels of diversity in these populations and previous estimates of their effective population sizes [39].

Since the number of loci affected by global selection is large, in chromosomal regions with low recombination rates Hill-Robertson effects will tend to oppose global adaptation [40] *(Supplementary Text)*. Simulations of a non-recombining population initialized at the predicted mutation-selection balance evolved to lower fitness (Fig. S15), while recombination enabled genome composition to be efficiently maintained under global selection (Fig. S14). Similar behavior in a different simulation model has previously been shown [40]. Consistent with this prediction, we found that chromosomes whose word frequencies exhibit deviations from the genome-wide average tend to be non-recombining sex chromosomes or to have smaller genetic length than average (Fig. S10). In the human genome, the genome-wide average number of mutations per crossover per generation is 0.87 [41], while in bacteria homologous recombination replaces small fragments of a genome with homologous fragments from other cells, and occurs with rates per site that are comparable to or greater than *u* [42], allowing global selection to maintain genome motif composition.

## 3 Discussion

We presented several lines of evidence indicating that genome-wide word frequencies have been shaped by global selection over long evolutionary timescales. First, we showed that extensive *in vitro* binding data exhibit statistically significant correlations with genome-wide relative word frequencies. These correlations are usually negative, indicating that strength of selection against a word scales with binding strength. Second, we identified a general hallmark of global selection - the strongly correlated frequencies of similar words - which we observed in all genomes. We argued that this correlation results from the biophysics of protein-DNA binding: since similar words have similar binding strengths, the evolutionary pressures they experience should likewise be correlated, resulting in the observed word-neighbor correlations. Third, we analyzed a wide range of null models, which demonstrated that word-neighbor correlations cannot be attributed to mutational biases. Fourth, we introduced an evolutionary model that was used to infer the global selective coefficients of words. Fifth, we analyzed the phylogenetic signal of global selection, which we now discuss.

The persistence of a phylogenetic signal in relative word frequencies over the large evolutionary distances seen in Figs. 5, S17, & S18 cannot be explained by a neutral divergence process. For example, *D. melanogaster* and *M. musculus* diverged over 600 Mya, yet their relative word frequencies (2-mer null) exhibit a correlation of *ρ* = 0.43 (Table S4). Billions of generations separate these genomes, and considering their per bp per generation mutation rates are ~ 10-^8^ [43] on average every neutral position will have mutated one or more times. Since our analysis was performed in introns, most of the ancient signal would have been destroyed leaving essentially no correlation. Furthermore, we showed that random drift does not play a major role in determining word frequencies by separately analyzing the chromosomes within each genome, which would drift independently of each other but instead cluster together in the PCA and exhibit strong correlations (Table S4). We propose that this phylogenetic signal persists because DNA binding motifs are conserved over long evolutionary timescales, and they continue to exert similar global pressures on genomes. While *cis*-regulatory architectures can evolve relatively rapidly, the motif specificities of homologous transcription factors from distant species have been shown *in vitro* to be highly conserved and evolve on a much slower timescale [44].

Our mathematical model, which represents global selection at mutation-selection balance, is appropriate for describing the maintenance of genome-wide word composition over evolutionary timescales. We find the selective coefficients of most words span within one order of magnitude of the point mutation rate *u* (Fig. 4), and since *u* ~ 10^−10^ per generation (*E.coli* [33]) or ~ 10^−8^ (humans [37]), we suspect these are among the smallest selective effects that have pervasively contributed to evolution. The model, however, cannot be used to infer the magnitude of selection that has acted periodically to change global composition. These potentially much larger effects – which we speculate would occur over timescales comparable to speciation events due to evolution of TFs with new binding motifs or substantial changes in expression levels of DNA-binding proteins – could be probed by a detailed comparative analysis of lineages that are at different stages of speciation. Given the key role of recombination in this process, further insights into global adaptation rates may result from detailed modeling of the interaction of recombination with global selection in combination with genomic analyses. Here, we showed that without recombination, genomes are only able to partially adapt globally, and cannot maintain genome compositions that have significantly higher global fitness (Figs. S14 & S15), a result which is consistent with analysis of chromosome sizes as well as Y chromosomes (Fig. S10). The presence of global selection should therefore result in a selective advantage for recombination [40].

Our extensive analysis of genomic data demonstrates that global selection is a universal evolutionary force that acts on genomes. We propose that this force arises from the functions of a diverse and distinct set of DNA binding processes within each genome that together generate a characteristic set of global pressures. As global selection maintains genome-wide motif compositions over long evolutionary timescales, its hallmark can be detected in the correlated frequencies of words and their mutational neighbors. Our findings introduce a new view on genomic evolution, in which molecular diversity that is effectively neutral over shorter timescales provides the raw material for global selection acting over much longer timescales.

## Materials and Methods

All methods are provided in *Supplementary Methods*.

## Supplementary Information

Includes: Methods (1), Text (1), Figures (S1-S23), Tables (S1-S5), and Data File (S1).

## Acknowledgements

We wish to thank B. Greenbaum, A. Grosberg, O. Hallatschek, E. Koonin, S. Leibler, B. Levin, M. Lynch, D. Pollock, M. Purugannan, M. Rockman, D. Roos, I. Ruvinsky, E. Shakhnovich, B. Shraiman, S. Small, and J. Weitz for valuable discussions. This work was supported by a Human Frontiers Science Program Young Investigator’s Grant.

## 1 Materials and Methods

### 1.1 Genomic sequence datasets

Sequences were downloaded from: ftp.ncbi.nlm.nih.gov/genomes/. The list of genomes for bacteria and archaea is found in the directory /GENOME REPORTS/prokaryotes.txt, and for eukaryotes in the directory /GENOME REPORTS/eukaryotes.txt. Bacterial plasmid sequences were not used. We manually curated a large, representative set of genomes across all bacterial and archaeal groups (947 genomes). For eukaryotes, all fully assembled and annotated genomes were included in the intron analysis (75 species, 1,304 chromosomes; mitochondria and plastid sequences were not included; see Table S3). Among eukaryotes, exon analysis was performed for 17 species (251 chromosomes; see Table S3). To extract the intron segments, CDS coordinates were obtained from the annotation files, and then mapped to the full length genome. For both exon and intron extraction, only one of the splicing isoforms was selected at random.

### 1.2 Measuring genomic word statistics

Word frequencies were collected by a sliding window of *k*-bp (*k* = 2,…, 8) across the sequences. For the coding region analysis, frequencies were collected on each open reading frame (i.e. multiple exons were joined for eukaryotes) and then combined for all open reading frames on a chromosome. Start and stop codons were ignored. For the intron regions, each intron segment was read separately and then the frequencies were combined. Gaps (N tracts in the eukaryotic sequences), if encountered, were replaced randomly by four bases at equal frequencies. Word frequencies were counted on the plus strand of DNA, and the counts of reverse-complement words were combined.

### 1.3 Constructing the null models in exon and intron sequences

For the coding region analysis using codon shuffling as the null model, synonymous codons were shuffled across the entire chromosome sequence. Start and stop codons were ignored. Each shuffled sequence was then scanned as above. The expected word frequencies 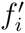 were calculated as the average of word counts from 1,000 such shuffled sequences. For the intron analysis using nucleotide base composition as the null model (1-mer null), base compositions were calculated on the chromosomal scale. For a word *w* with *N*_*A*_, *N*_*T*_, *N*_*C*_, *N*_*G*_ counts of A, T, C, and G bases, respectively, the expected frequencies were calculated as 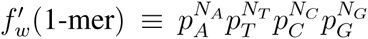, where *p*_*A*_, *p*_*T*_, *p*_*C*_, *p*_*G*_ are base compositions measured from the full length sequences. For analysis using the dinucleotide (2-mer null) or trinucleotide (3-mer null) models, a Markov chain model was used to compute expectations. A *k*-mer word *w* = (*w*_1_*w*_2_*w*_3_… *w*_*k*_) is composed of an overlapping set of dinucleotides *(w*_1_*w*_2_), (*w*_2_*w*_3_),…, *(w*_*k*-1_*w*_*k*_). The probability of observing each dinucleotide (ab) was measured across all introns conditional on a being at the first position, and denoted *p(b| a*). Using these measured values, the expected frequency of the word *w* is given by

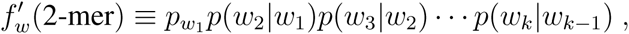

where *p*_*w*_1__ is the measured frequency of nucleotide *w*_1_. Similarly, for the trinucleotide model, the word is composed of an overlapping set of trinucleotides, and

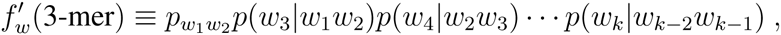

where *p(c| ab*) is the probability of observing trinucleotide (*abc*) conditional on the dinucleotide (*ab*) occupying the first two positions; and *p*_*w*_1_*w*_2__ is the measured frequency of dinucleotide (*w*_1_*w*_2_).

### 1.4 Analysis of variance (ANOVA) of genomic word frequencies

To determine the extent to which the dinucleotide and trinucleotide composition of genomes can explain *k*-mer frequencies, we performed a multi-factor ANOVA of word frequencies vs. their dinucleotide and trinucleotide composition. The raw frequencies of 6-mer and 8-mer words were measured in introns, and their relative frequencies were computed under the 1-mer, 2-mer, and 3-mer null models. For the dinucleotide ANOVA, the composition of each word was recorded as a 10-dimensional vector, indicating the counts of each of the following dinucleotides, identifying reverse-complement pairs as a single factor: (AC/GT, CA/TG, AT, TA, CT/AG, TC/GA, CG, GC, AA/TT, CC/GG). For the trinucleotide ANOVA, the composition of each word was given by a 32-dimensional vector, indicating the counts of each possible trinucleotide, identifying reverse-complement pairs as a single factor.

Since dinucleotides and trinucleotides overlap within any *k*-mer word (*k* ≥ 4), there exists a relation among the vector components which causes a singularity in performing regression analysis or ANOVA. A procedure to remove such degeneracies is known as differencing, in which two components are replaced by a single component that measures their difference. For dinucleotide analysis, we replaced the components corresponding to AT and TA by a single component that measures the difference in counts of these two dinucleotides. For trinucleotide analysis, we replaced the components corresponding to CGA and TAA by a single component that measures the difference in counts of these two trinucleotides. The resulting regression is then non-singular, and we verified that the results were not sensitive to the choice of differencing components.

Using the dinucleotide analysis, we also measured the fraction of the total variance explained by the CpG mutational neighborhood, which consists of four dinucleotides: CG, CT/AG, CA/TG, CC/GG. This was computed by performing the ANOVA using the four factors in the CpG mutational neighborhood.

### 1.5 Principle components analysis of genomic word frequencies

Principle component analyses were performed on the correlation matrix between relative word frequencies using the 2-mer null model. For eukaryotic intron regions (Figs. 5, S17, S18), relative word frequencies of 1,049 chromosomes from 61 species were used for the PCA (Fig. S17 legend and Table S3). Excluded were chromosomes that were shorter than 1 Mb or having < 0.8 correlation between the measured relative frequencies or excess pressures of reverse complement words on the plus strand, as well as the chromosomes of the bird *Ficedula albicollis* because its word frequencies take on a split distribution in the 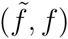 plane. The word frequencies on these chromosomes were so extreme that they dominated the PCA analysis. The plots for subgroups shown in each panel were produced by PCA on each subgroup alone. For bacterial coding regions (Fig. S19 & S20), relative word frequencies were computed using codon shuffling as the null model, and counts were combined on both strands of all analyzed exon sequences.

### 1.6 Simulation of global selection

In each simulation the population consisted of *N* word composition vectors *f*^(1)^,…, *f*^(*N*)^, with 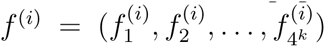 giving the *k*-mer word counts in sequence *i*, and 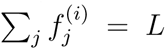. The populations were initiated from random sequences by sampling *N* times from a multinomial distribution Mult(*f*^*rand*^, *L)* with *f*^*rand*^ = (1/4^*k*^,…, 1/4^*k*^). Populations were propagated through rounds of Wright-Fisher reproduction, with each round consisting of (1) mutation, (2) recombination, and (3) selection. During the mutation step, for each sequence *i*, the total number of mutated words was sampled from a Poisson distribution with mean *kuL*. Each mutation caused a jump from word *w* to word *w’*, where *w* was sampled based on the composition *f*^(*i*)^, and *w’* was sampled uniformly from the mutational neighbors of *w*. With each jump, *f*^(*i*)^ is updated, and the next mutation was sampled based on the updated *f*^(*i*)^. During recombination random mating was implemented in the population, and for each mating pair *i* and *j* a fixed fraction of the sequence *L* corresponding to a total of l words was exchanged between *f*^(*i*)^ and *f*^(*j*)^: *f*^(*i*)^ → *f*^(*i*)^ + ∆*f*^*(j →i)*^ - ∆*f*^*(i→j)*^ and *f*^*(j)*^ → *f*^(*j*)^ + ∆*f*^(*i→j*)^ - ∆*f*^(*j→i*)^, where ∆*f*^(*j→i*)^ and ∆*f*^(*i→j*)^ were sampled from Mult(*f*^(*j*)^, *l*) and Mult(*f*^(*i*)^, *l*), respectively. During selection, the fitness of each sequence *i* was calculated as *e*^*s·f*^*(i)*^^, and we define *p*_*i*_ = *e*^*s·f*^*(i)*^^ / Σ_*j*_*e*^*s·f*^*(j)*^^ to be the probability distribution for selecting sequences to populate the next generation. The next generation of composition vectors is then obtained by a single sampling from Mult(*p*, *N)*. To introduce bottlenecks, all steps were the same except that during population size transitions, from *N*_*large*_ → *N*_*small*_, or from *N*_*small*_ → *N*_*large*_, the sequences for the next generation were selected randomly from Mult(*p*, *N*_*small*_) and Mult(*p*, *N*_*large*_), respectively.

## 2 Supplementary Text

### 2.1 Mathematical Model of Global Selection

We use a generalization of the well-known quasi-species model [1], applied to the evolution of genome-wide word frequencies. We summarize the basic formulation of the model, which follows from our previous work [2], and then show how to invert the solution to obtain the exact mapping of observed frequencies to selective coefficients, which is used throughout the main text. Lastly, we discuss the simulation results on finite population sizes, Muller’s ratchet, recombination, and bottlenecks.

#### 2.1.1 Model derivation

A sequence of length *L* is represented by a *k*-mer composition vector f in the projected space 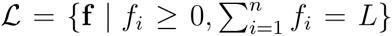, where *n* = 4^*k*^ is the total number of possible words, and *f*_*i*_ is the number of occurrences of word *i* in the sequence. We denote by s the selection vector whose *i-*th component is the selective coefficient *s*_*i*_ acting on each word of type *i*. We consider a large population of evolving sequences, each represented by a composition vector. At each generation, each sequence contributes offspring proportional to its total fitness, exp(s · f). The offspring mutate according to a mutational transfer operator, *G*(**f | f’**), which gives the probability of a mutational transition **f’ → f**. Under these dynamics, the expected frequency of f within the population at generation *t*, denoted by *P*_*t*_(**f**), evolves according to the equation

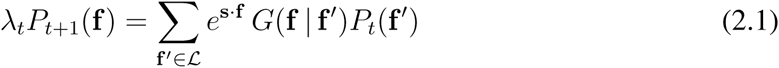

where λ_*t*_ is a normalizing factor, which measures the average population fitness at generation *t*, given by summing both sides over all values of f. The irreducibility of the transfer operator *G* guarantees (by the Perron-Frobenius theorem) that the linear mapping of *P*_*t*_ → *P*_*t*+1_ converges to a unique steady-state distribution, *P*(f), which dictates the population structure at mutation-selection balance.

Since each locus along DNA mutates independently, the transfer operator can be decomposed as a product over all loci, which implies that *P*(f) is a multinomial distribution [2]. A multinomial random variable x, denoted x ~ Mult(p, *L*), results when sampling *L* independent events from a discrete probability distribution *p*_*i*_, where *x*_*i*_ is the number of outcomes of type *i*. With this notation, we have f ~ Mult(p, *L*), where p is the single-locus distribution of words at mutation-selection balance. This distribution p is obtained as the unique positive solution of the eigenvalue problem

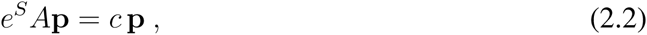

where *S* = diag(s), and *A*_*ij*_ is the probability that word *j* mutates into word *i*. Once more the Perron-Frobenius theorem guarantees a unique positive eigenvector p with associated eigenvalue c, which determines the equilibrium average fitness, λ_*t*_ → λ = *c*^*L*^.

The above eigenvalue problem can be solved numerically, e.g. to obtain the plots in Fig. 2. To invert the relationship, and obtain an expression for *s*_*i*_ given p, we use the fact that the point mutation rate *u* (per bp per generation) is extremely small. We express A explicitly as *A* = *e*^*uk(M-I)*^, where *I* is the identity matrix and *M*_*ij*_ is the probability that a single mutation converts word *j* into word *i*; *M*_*ij*_ is non-zero only for words *i* and *j* that are mutational neighbors. In the following analysis, we assume unbiased mutation, i.e. *M*_*ij*_ = 1/(3*k*) for any pair of mutational neighbors *i* and *j*; generalization for biased mutations is straightforward, discussed below. We expand *A* in small *u* to first order, i.e. neglecting contributions from double (or higher order) mutants that occur within a single word locus. Subsituting *A* ≈ *I + uk(M - I)* in (2.2) yields,

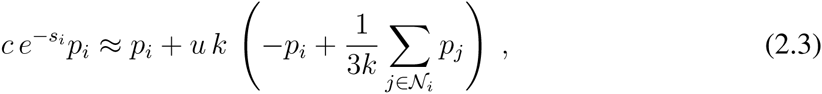

where 𝒩_*i*_ is the set of mutational neighbors of word *i*. The second term within the parentheses is 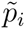, the average frequency of mutational neighbors. Dividing both sides by *p*_*i*_ and taking logarithms, we find

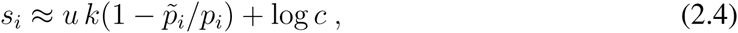

which after division by *u* yields equation (1) of the main text. This expression shows that the values of *s*_*i*_ are determined by 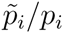 up to an additive constant. In the main text, pressure distributions are shown using a value of *c* = 1, which corresponds to taking the average global fitness of a sequence to be zero.

Known mutational biases *M*_*ij*_ can easily be incorporated in the above, by defining 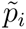 to be a weighted average of the mutational neighbors:

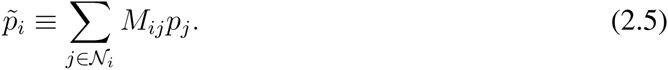

Since our goal was to survey a large number of species across all domains of life, most of which have not been characterized as far as mutational biases, we developed the excess pressure method, which correctly infers selective coefficients when mutational biases are not known (see next section).

We note that our mathematical model ignores the local, overlapping structure of words within a genome, effectively representing the genome as a ‘bag of words’. This approximation is fully justified because the operation of shifting a *k*-mer window by one position along a DNA sequence results typically in a jump to a very different word. The only exceptions occur within a very small subset of loci that contain short sequence repeats of size ≥ *k*.

#### 2.1.2 Finite Populations and Muller’s Ratchet

The above analysis is strictly valid only in the infinitely large population limit. For finite populations and values of *L ≫ 1*, deviations from this equilibrium can occur due to Muller’s ratchet [3, 4]. This result has been discussed in [5], and we briefly review the same reasoning here. Due to the large genotype space, the maximally fit sequence - i.e. one in which the most favorable word is present at each locus - is rapidly lost in the population, because the rate of mutation away from this sequence, *Lu*, is much larger than the rate of bac*k*-mutation, *u*. Within a few generations, due to the finite population size, every sequence has accumulated at least one deleterious word, and the fittest sequence in the population is now one which carries at least one mutation. This process drives the population away from the fittest sequence, and reaches an equilibrium distribution some distance from **P(f)**, which depends in a complex manner on *N, L*, and *u*[5]. Similarly, in genomic regions with low recombination rates, interference effects of various kinds (known as Hill-Robertson effects; see [6]) will oppose global adaptation. Recombination, however, provides an accessible route to reverse the ratchet effect, by enabling in each generation the previously fittest sequence to be recovered. Our simulations demonstrate that the equilibrium *P*(f) is indeed achieved in a finite size, recombining population (Figs. S14,S15).

### 2.2 Mutational Biases and Genomic Word Composition

In this section we analyze the role of mutational biases for the word frequency statistics of genomes. Direct measurements of mutational biases are performed by mutation accumulation experiments in which populations are propagated over many generations under conditions that minimize the strength of selection, typically by frequent passage through strong bottlenecks. Such studies have been performed in *E.coli* [7], *B. subtilis* [8], *S. cerevisiae* [9, 10], *C. elegans* [11], *D. melanogaster* [12], *A. thaliana* [13], and other species [14]. Additionally, measurements using known pedigrees have been made in humans [15, 16]. Mutational biases at the single nucleotide level are characterized by a mutational transition matrix among the basepairs (1-mers) AT, TA, GC, and CG, with six possible mutation types: AT → GC, GC → AT, AT → TA, GC → TA, AT → CG, and GC → CG. The rates of these mutation types vary within about one order of magnitude and differ between species. For example, in *H. sapiens* the rate of GC → AT is about 6 larger than AT → CG, while in *A. thaliana* it is about 14 times larger (see [15] Table 2). These rates are context independent, since they are measured at the single nucleotide level without considering neighboring nucleotides.

Context-dependent biases have also been measured in different species. The strongest known bias in vertebrates is due to CpG methylation, which increases the overall rate of mutation at CpG dinucleotides by about tenfold on average, and which varies across species [17]. In *E.coli*, different dinucleotide contexts were found to affect mutation rates, particularly for transitions [7]. In *B. subtilis*, trinucleotide contexts were assessed, and while dinucleotide contexts were found to account for most context dependencies, certain trinucleotide contexts were statistically significant [8]. In yeast, where mutation accumulation experiments have been the most extensive, measurements on trinucleotide contexts found that a small number of the 32 possible trinucleotide contexts explain most of the variance in mutation rates [9]. Among the 16 trinucleotide contexts for AT basepairs, no statistically significant differences were detected, while among the 16 contexts for GC basepairs two contexts (CCG/CGG and TCG/CGA) had elevated rates by a factor of two. Most of the rate variability could therefore be accounted for by two bits of data: the nucleotide identity of the central basepair (AT or CG), and whether or not the nucleotide is in either of CCG/CGG or TCG/CGA contexts.

#### 2.2.1 Word frequency statistics

It is well-known that the equilibrium frequency of A/T nucleotides predicted based on context-independent mutational biases deviates significantly from the observed A/T genome composition at silent sites in many species (see e.g. [18, 19, 15]). To determine whether context-dependent mutational biases can explain the *k*-mer statistics of genomes, we performed the two separate analyses described next.

First, we constructed Markov chain null models of genomic sequences in which the frequency of a nucleotide was conditional on the previous nucleotide (2-mer null) or the previous dinucleotide (3-mer null) (see Methods, Sec. 1.3). Under the null hypothesis that dinucleotide or trinucleotide context-dependent mutational biases account for observed word frequencies, the frequency of any *k*-mer with *k* > 3 will be determined by the conditional probabilities of its constituent 2-mers or 3-mers. Under this null hypothesis, for each genome we parameterized the 2-mer and 3-mer null models using all intron sequence, and computed the expected frequency 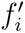 of each word *i*. For each analyzed genome, given total intron length *L*, we recorded the observed counts of each word *i* as *C*_*i,obs*_, and computed the expected counts of each word, 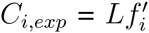, and its variance 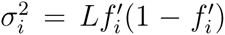 under each null model. We then computed the z-score of each word as *z*_*i*_ = (*C*_*i,obs*_ - *C*_*i,exp*_)/σ_*i*_. Given the large values of *L* for each genome, the distribution of z-scores is expected to be approximately normally distributed with mean zero and variance one. Instead, we observed that the distribution of z-scores spans 100’s of standard deviations under both null models, indicating that the statistics of higher *k*-mers cannot be explained by context-dependent mutational biases (Figs. S6A & S6B).

Second, we performed a multi-factor analysis of variance (ANOVA) of word frequencies against their dinucleotide or trinucleotide composition. This analysis was designed to measure the fraction (*r*^2^) of the total variance of word frequencies that could be explained by dinucleotide and trinucleotide composition, and hence by context-dependent mutational biases. Since the analysis is equivalent to regression of word frequencies on di-and tri-nucleotide factors, the *r*^2^ value measures the best possible fit of the model to the data, and should thus be considered an upper bound on its explanatory power. For example, in the trinucleotide null model, the regression involves 32 factors, but in reality we know that far fewer trinucleotide contexts exhibit significant mutational biases (see above). The regression model will therefore over-fit the data, hence the calculated *r*^2^ will be higher than its true explanatory capacity.

The ANOVA results are given in Table S5, performed using raw or relative word frequencies (either 6-mer or 8-mer) vs. the dinucleotide or trinucleotide factors. The ANOVA on 8-mer frequencies relative to the genome nucleotide composition (indicated in the *f* (1-mer) column), shows that for mouse and human genomes, respectively 34% and 29% of the variance is accounted for by dinucleotide composition, while these numbers increase only slightly to 39% and 33% for trinucleotide composition. This leaves more than 60% of the variance unexplained by these compositional models. Normalizing 8-mer frequencies relative to the 2-mer null model, and performing the ANOVA on trinucleotides (shown in the *f* (2-mer) column), we find the trinucleotide composition explains only 1% and 8% of relative word frequencies in mouse and human genomes, respectively. Thus, the increase of model complexity from dinucleotides to trinucleotides adds 22 degrees of freedom in regression analysis, but results in little additional explanatory power. In both of these genomes, the CpG mutational neighborhood (i.e. dinucleotides within 1 mutation of CpG) accounts for most of the dinucleotide signal, explaining 32% of the variance in mouse and 26% in human (see *f* (1-mer) column). Explained variance exhibits a wide range across genomes, where dinucleotides explain as little as 15% of the variance in *C.elegans* to as much as 55% in *G. gallus*, with similar trends observed for trinucleotides. A larger proportion of the variance can be explained for 6-mer frequencies, which is likely due to the much larger number of 8-mers (32,896) vs. 6-mers (2,080), since over-fitting by the regression becomes more difficult for the 8-mer data.

The ANOVA analysis also provides a simple consistency check on the Markov chain null models described above. The very small *r*^2^ values given in the *f* (2-mer) column of the dinucleotide ANOVA indicate that normalization by expected frequencies computed according to the Markov chain dinucleotide model (Methods, Sec. 1.3) effectively accounts for most of the dinucleotide signal in the word frequency data. And, similarly, for the *f* (3-mer) column of the trinucleotide ANOVA.

We conclude that context-dependent mutational biases are unable to explain the wide distribution of word frequencies we observed. The fraction of the variance that is attributable to mutational biases is mainly due to dinucleotide biases, and largely dominated by the CpG mutational neighborhood in vertebrates and plants. For this reason, we used the 2-mer null model when presenting results on selection coefficients and PCA in the main text.

#### 2.2.2 Word-neighbor correlations

The word-neighbor correlations we detected in all genomes were calculated using five different null models in several different genomic regions: in exons, synonymous codon shuffling (Figs. 2A, S3, S4, S7), synonymous dicodon shuffling (Fig. S23); in introns, 1-mer null (Fig. S5), 2-mer null (Fig. S6A), and 3-mer null (Fig. S6B); in repeat-masked intronic regions, 1-mer and 2-mer null (Fig. S21); and in DNAse I hypersensitive regions, 1-mer and 2-mer null (Fig. S22). The ANOVA results above show that our normalization by 2-mer and 3-mer expected frequencies effectively removes correlation with dinucleotide and trinucleotide composition. To separately verify that the word-neighbor correlations are not accounted for by 2-mer and 3-mer composition, we regressed out their compositional effects and examined word-neighbor correlations of the residuals (Fig. S16). The strongly correlated residuals we observed provide further evidence that word-neighbor correlations do not result from context-dependent mutational biases.

In bacteria, where genomes are dominated by exon regions, our most stringent test is the synonymous dicodon shuffling scheme, which swaps synonymous codons while preserving the coding sequence as well as the genome’s dicodon composition. This means that the compositional biases on all *k*-mers for *k ≤* 4 are preserved. Any remaining signal therefore cannot be attributed to context-dependent mutational biases. In Fig. S23, we see that highly significant word-neighbor correlations persist in word frequencies normalized according to dicodon shuffling statistics. It is important to note that this shuffling scheme is overly conservative, and it constrains sequences much more than they would be in nature. Despite this fact, the excess pressures inferred based on the dicodon shuffling statistics (Fig. S23) span a range of width about *5u - 10u* in the bacteria we examined indicating that lots of global pressures exist on word with *k ≥* 5 which cannot be adequately captured by the word statistics for *k ≤* 4.

#### 2.2.3 Excess pressures, mutational biases, and global selective coefficients

Since mutational biases have been measured in only a small number of species, we developed a method that can accurately infer global selective coefficients without requiring measurements of mutational biases as input. We therefore defined the *excess pressure* to be the selective pressure required to shift the word frequencies away from their expectation based on a given null model. To do this we determine a set of effective selective coefficients 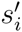 that would maintain the genome’s statistics at the expectation based on the given null model. We use an unbiased transition matrix and infer the values 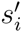 from the expected frequencies 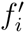 using Eq. 1. Next, we calculate the selective coefficients *s*_*i*_ needed to maintain the genome’s actual statistics at the observed word frequencies *f*_*i*_, again for an unbiased transition matrix, i.e. using Eq. 1. The excess pressures ∆*s*_*i*_ are defined as the difference 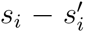. In this way, unknown mutational biases are represented using effective selective coefficients 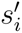, whose contribution is subtracted off from the measured values *s*_*i*_.

Using genomes where mutational biases have been measured, we tested the excess pressure method. We used the direct measurements to yield the mutational transition matrix *M*_*ij*_ defined in Sec. 2.1.1. Using this matrix in Eq. 2.5, we then computed selective coefficients 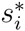 using Eq. 2.4. Since the transition matrix explicitly accounts for mutational biases, 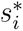 are the selective pressures required to shift the word frequencies away from the equilibrium established by mutation alone. In Fig. S12, we compared excess pressures *∆s*_*i*_ versus coefficients 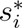, finding excellent agreement between the two methods. In human, plant, and fly genomes, the two methods exhibited correlations between 0.92 and 0.99. We conclude that excess pressures accurately assess global selective coefficients, and are thus useful when direct measurements of mutational biases are not available. Since most mutational biases are accounted for by dinucleotide effects, we expect that excess pressures computed in introns using the 2-mer null model (Figs. 4A, S6A) provide our most reliable estimate of global selective coefficients in eukaryotic genomes.

## 3 Legends

### 3.1 Supplemental Figures

**Figure S1:**
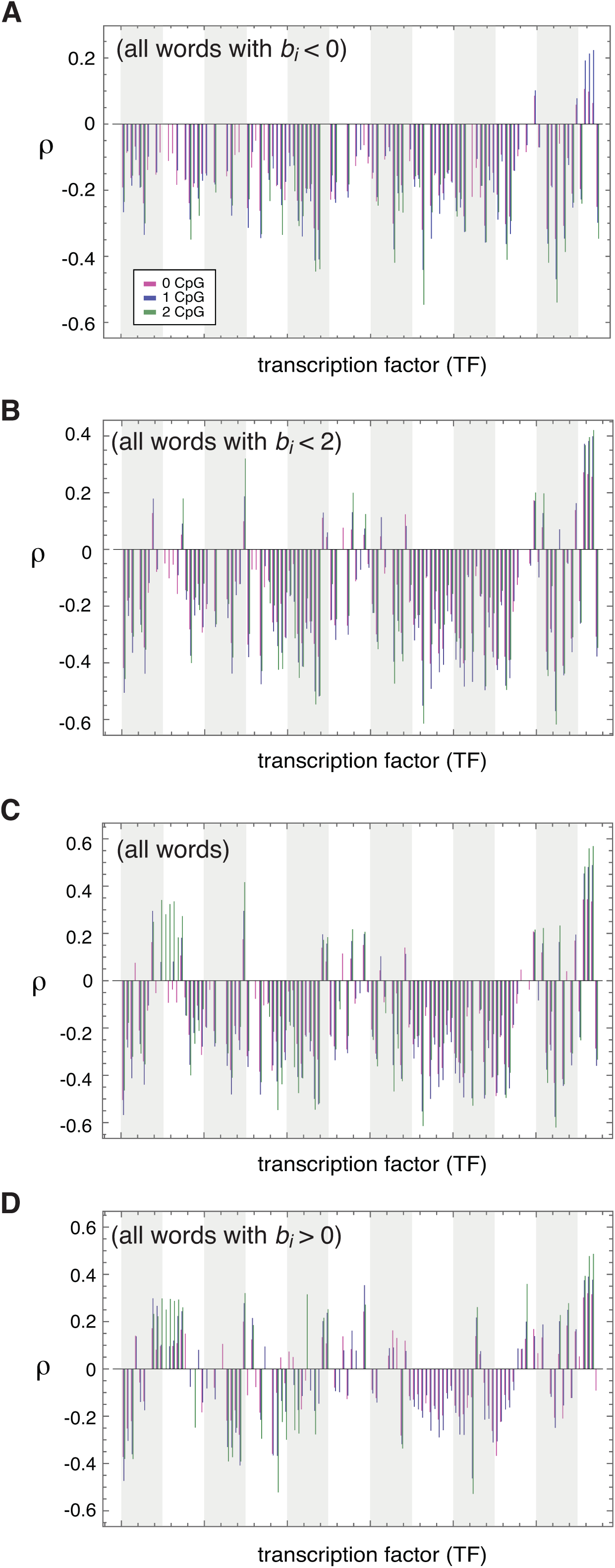
Mouse TF binding score correlation with genomic word frequency data. Correlation coefficients (Spearman’s ρ) computed between relative frequencies *f*_*i*_ and binding scores *b*_*i*_ for each mouse TF, where *i* range over all possible 8-mers. The binding scores used here corresponded to the z-scores reported in [20]. Frequencies *f*_*i*_ corresponded to the raw counts divided by expected counts based on nucleotide composition (i.e. the 1-mer null). Correlations are separately computed over subsets of words with the same CpG content. Bars are shown only for statistically significant correlations, with p-value < 10^−6^. Vertical gray and white blocks span 10 TFs each. Panels (A) and (B) use weak binding words, *b*_*i*_ < 0 or *b*_*i*_ < 2, respectively. Panel (C) does not condition on binding strength, while panel (d) uses words with *b*_*i*_ > 0.

**Figure S2:**
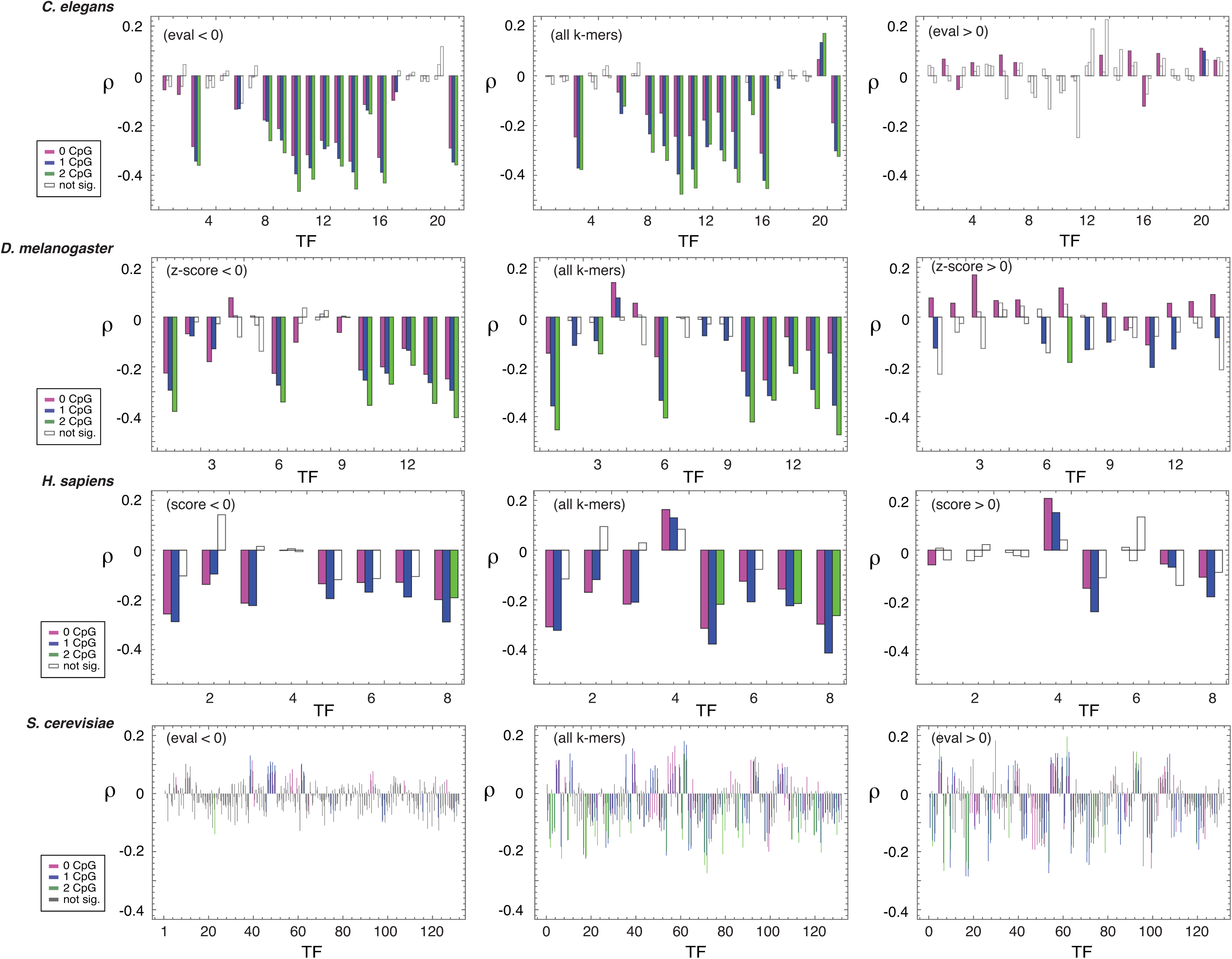
TF binding score correlation with genomic word frequency data for additional species. For each species and each TF, we computed the Spearman correlation (ρ) of binding scores *b*_*i*_ vs. genomic word frequencies *f*_*i*_, where *i* indexes over all 8-mer words. We plot ρ values on three different subsets of *k*-mer words: (left panels) using all words i satisfying *b*_*i*_ < 0; (middle panels) using all words; and (right panels) using words satisfying *b*_*i*_ > 0. Correlation values are either plotted in color bars when statistically significant (*p*-val < 10^−6^) or as white or gray bars otherwise. Negative correlations are observed for the majority of TFs in all cases, and occur most frequently over *k*-mers with *b*_*i*_ < 0 in mouse (Fig. S1), worm, fly, and human, and over *k*-mers with *b*_*i*_ > 0 in yeast. Frequencies *f*_*i*_ corresponded to the raw counts divided by expected counts based on nucleotide composition (i.e. the 1-mer null) for worm, fly, and human; counts were determined using all intron data in these species. In yeast, which has few introns, the raw counts were made across all exons, and the expected counts were obtained from the synonymous codon shuffling scheme. Binding scores *b*_*i*_ corresponded to z-scores for the fly dataset [21, 22, 23], and to e-values for worm [24] and yeast [25] datasets; the human dataset [26, 22] had 5 TFs with z-score data and 3 TFs with e-value data (see Table S1).

**Figure S3:**
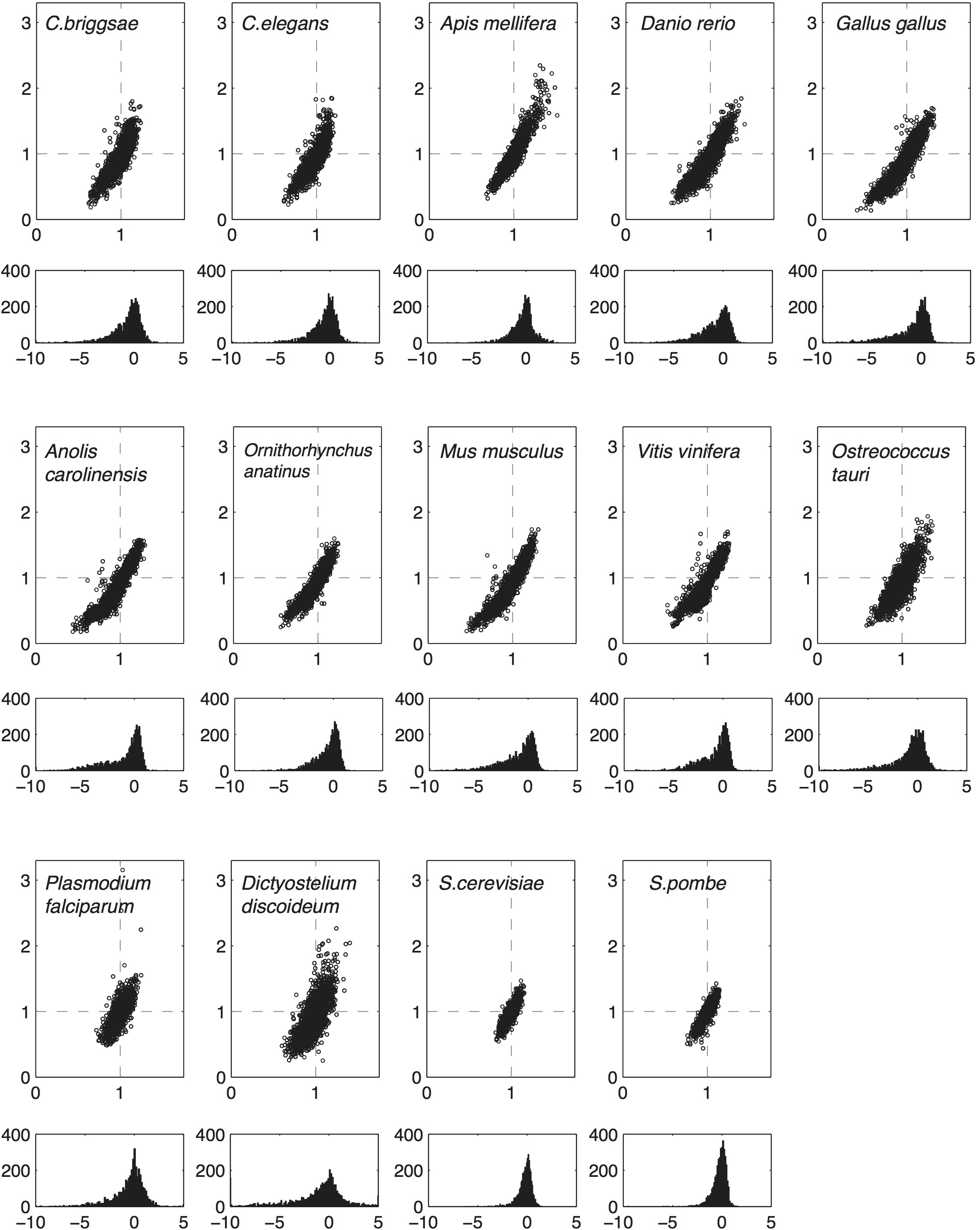
Word frequency plots and pressure distributions of exons for a range of eukaryotic species (*k* = 6). For each species, the top panel shows the 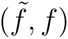 plot, and the bottom panel gives the corresponding excess pressure distribution, with ∆*s*_*i*_/*u* on the *x*-axis.

**Figure S4:**
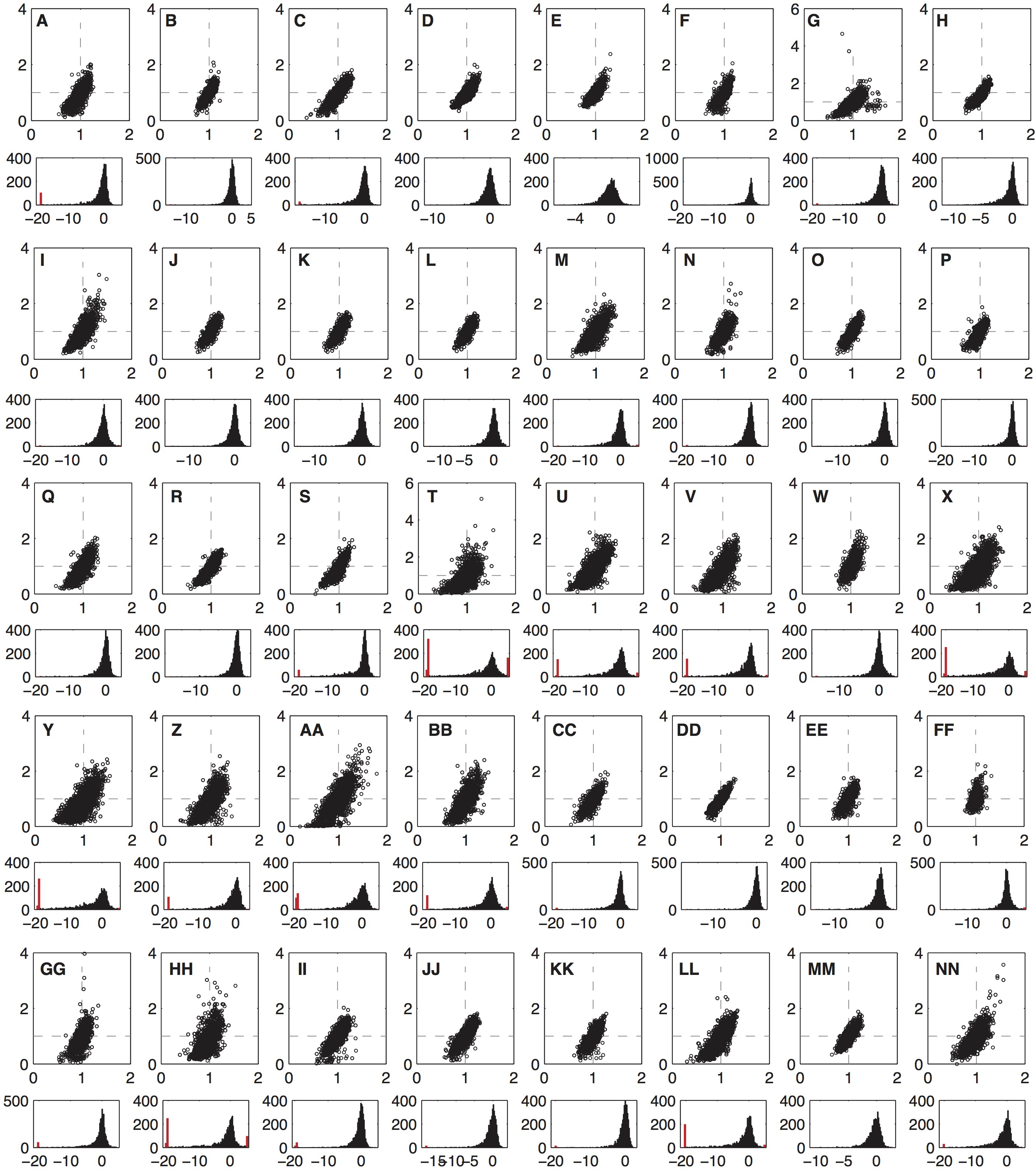
Word frequency plots and pressure distributions of exons for a range of bacterial species (*k* = 6). See Table S2 for species names, classification, and strain details. For each species, the top panel shows the 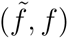 plot, and the bottom panel gives the excess pressure distribution, with ∆*s*_*i*_/*u* on the *x*-axis. Because certain bacteria exhibit a very wide pressure distribution, typically due to a small number of words with large negative pressures, we plot using red bars (3 on the left edge, 1 on the right edge) word counts with pressures in the ranges (-1000, −500), (-500,-100), (-100, −20), and > 5, respectively.

**Figure S5:**
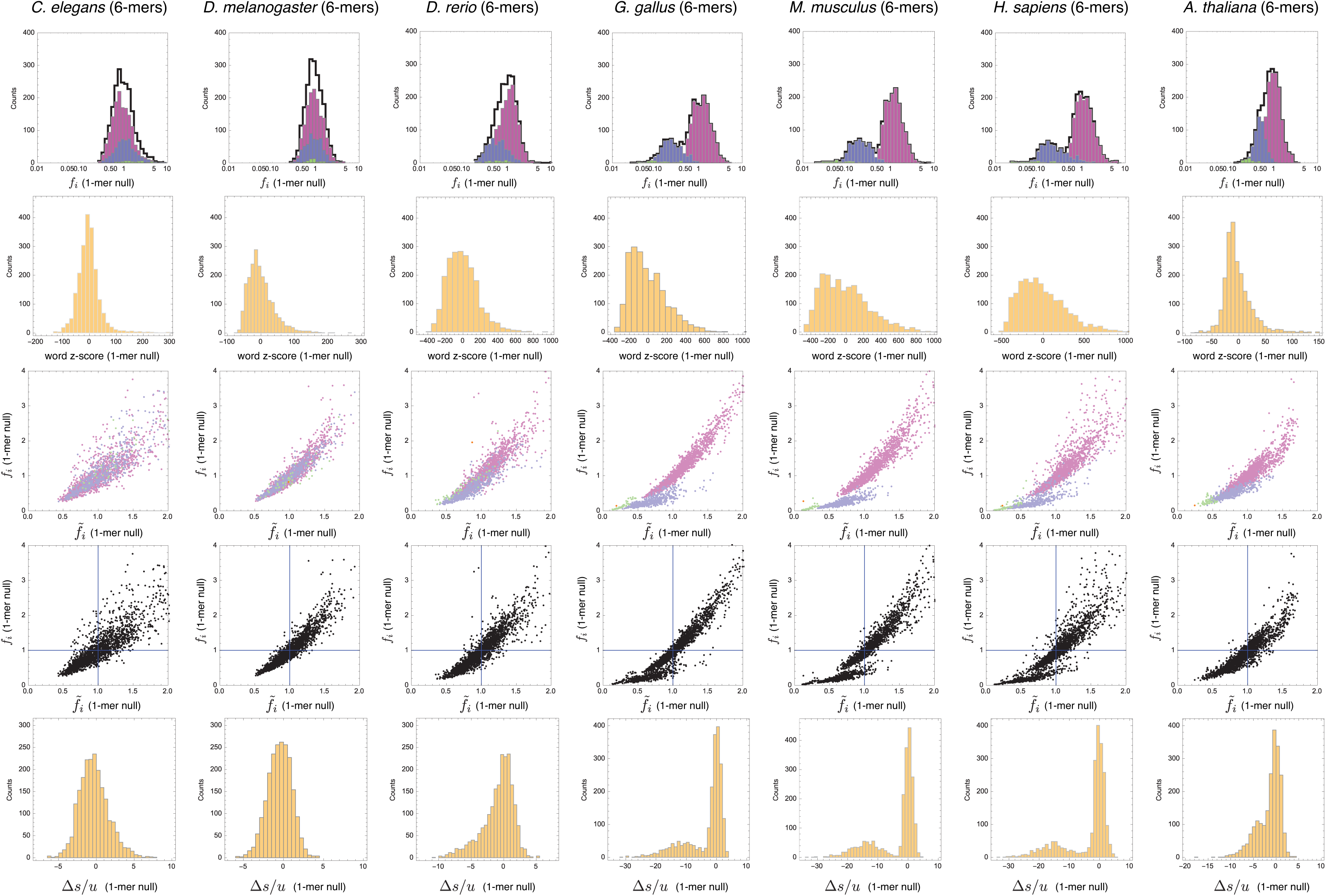
Analysis of intron word frequency distributions using the 1-mer null model. Word frequencies (*k* = 6) are shown relative to expectation based on nucleotide composition of introns (1-mer null). **Row 1**. Relative word frequency histograms. Black curve indicates all words, purple bars (0 CpG), blue bars (1 CpG), green bars (2 CpG’s). **Row 2**. Distribution of genomic word frequency z-score distributions. For each word *i*, its observed counts *C*_*i,obs*_ were measured, and the expected counts *C*_*i,exp*_ as well as expected variance 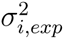 were computed using the 1-mer null. The word z-scores were computed as (*C*_*i,obs*_ - *C*_*i,exp*_)/*σ*_*i,exp*_. **Rows 3 & 4**. The 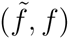 plot for each species; colors in Row 3 indicate CpG content as in Row 1. **Row 5**. The distribution of excess pressures relative to the 1-mer null.

**Figure S6A:**
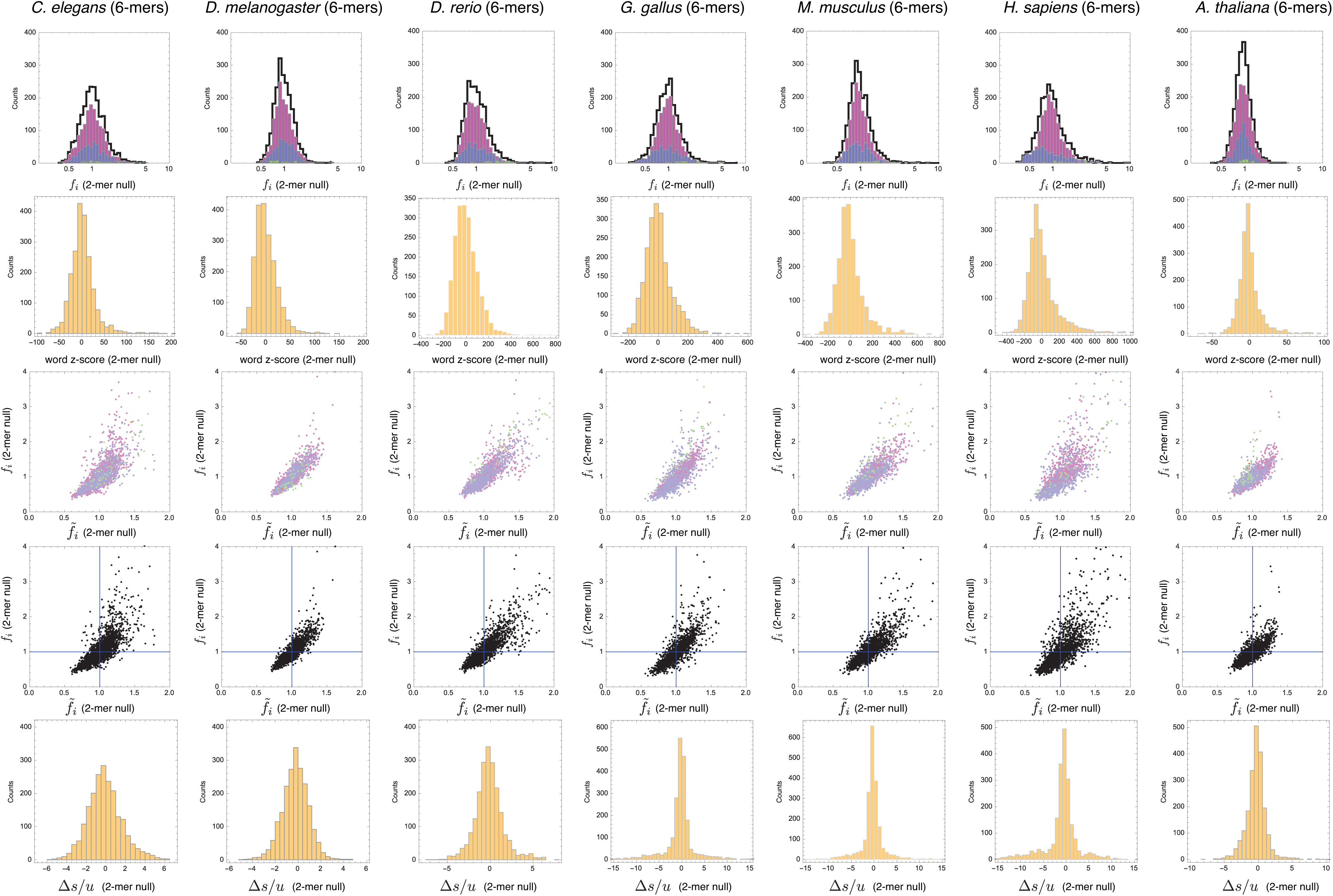
Analysis of intron word frequency distributions using the 2-mer null model. Caption identical to S5, except all word frequencies are computed relative to the dinucleotide composition using the Markov chain model described in Methods.

**Figure S6B:**
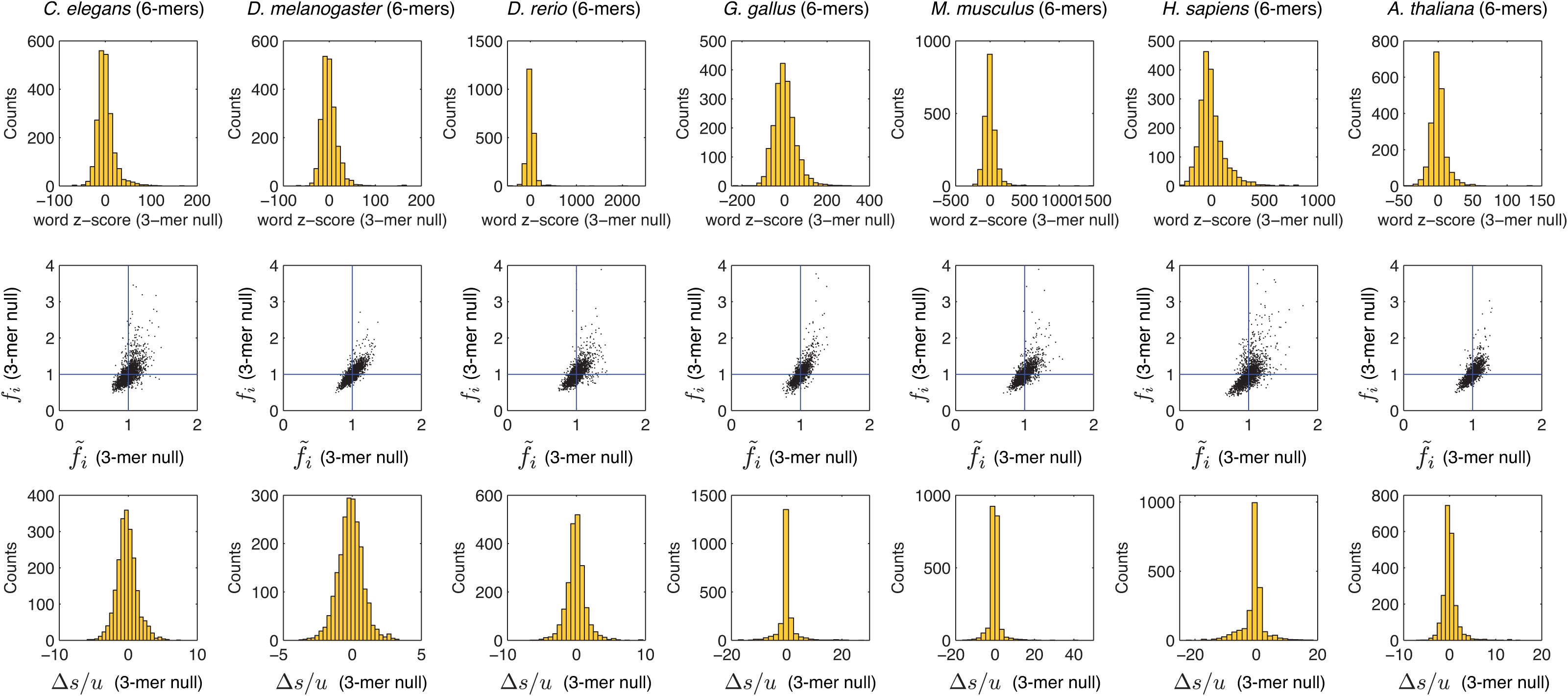
Analysis of intron word frequency distributions using the 3-mer null model. Caption identical to S5, except all word frequencies are computed relative to the trinucleotide composition using the Markov chain model described in Methods.

**Figure S7:**
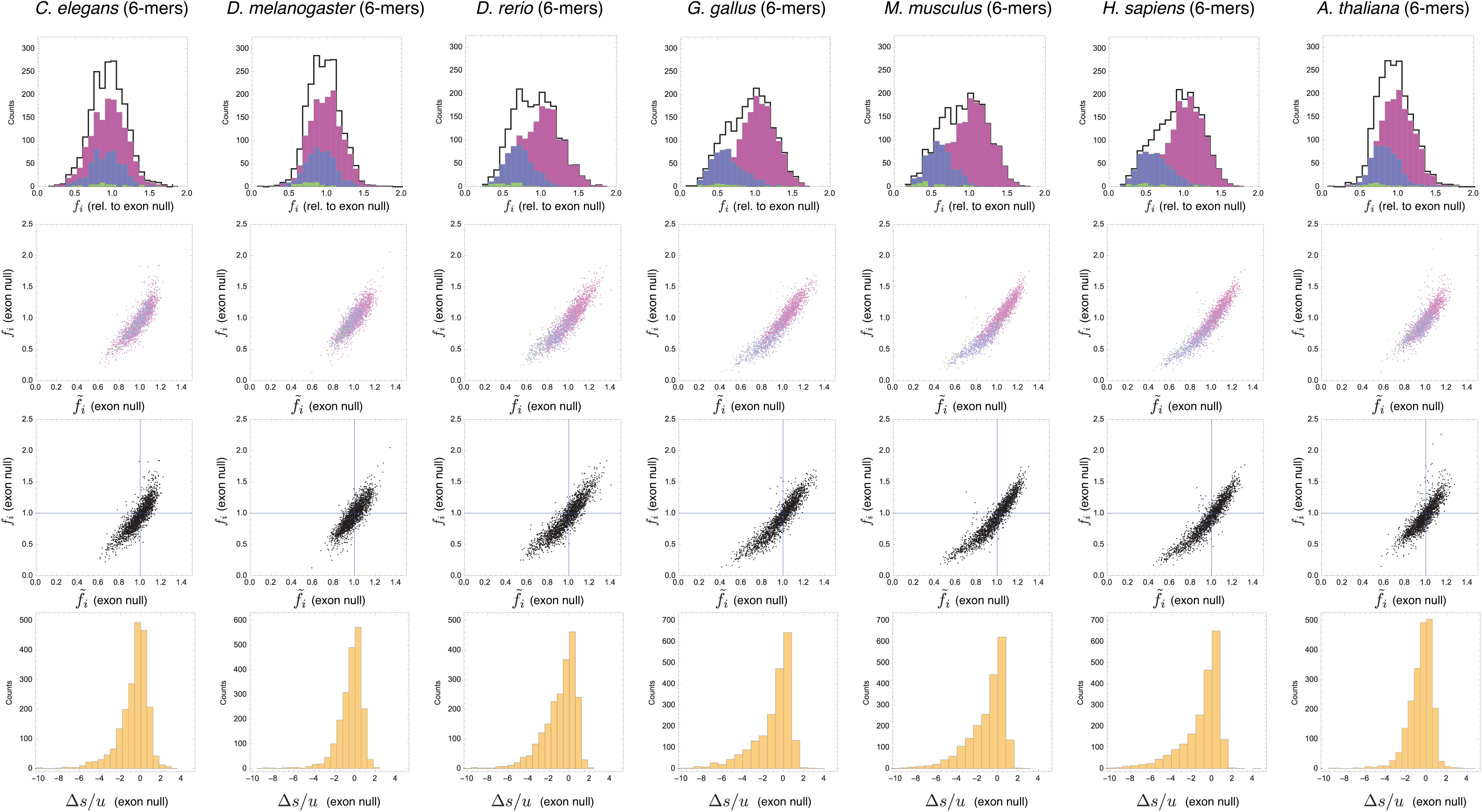
Analysis of exon word frequency distributions using the codon shuffling model. Word frequencies (*k* = 6) are shown relative to expectation based on the synonymous codon shuffling scheme in exons (exon null). **Row 1**. Relative word frequency histograms. Black curve indicates all words, purple bars (0 CpG), blue bars (1 CpG), green bars (2 CpG’s). **Rows 2 & 3**. The 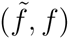 plot for each species; colors in Row 2 indicate CpG content as in Row 1. **Row 4**. The distribution of excess pressures relative to the exon null.

**Figure S8:**
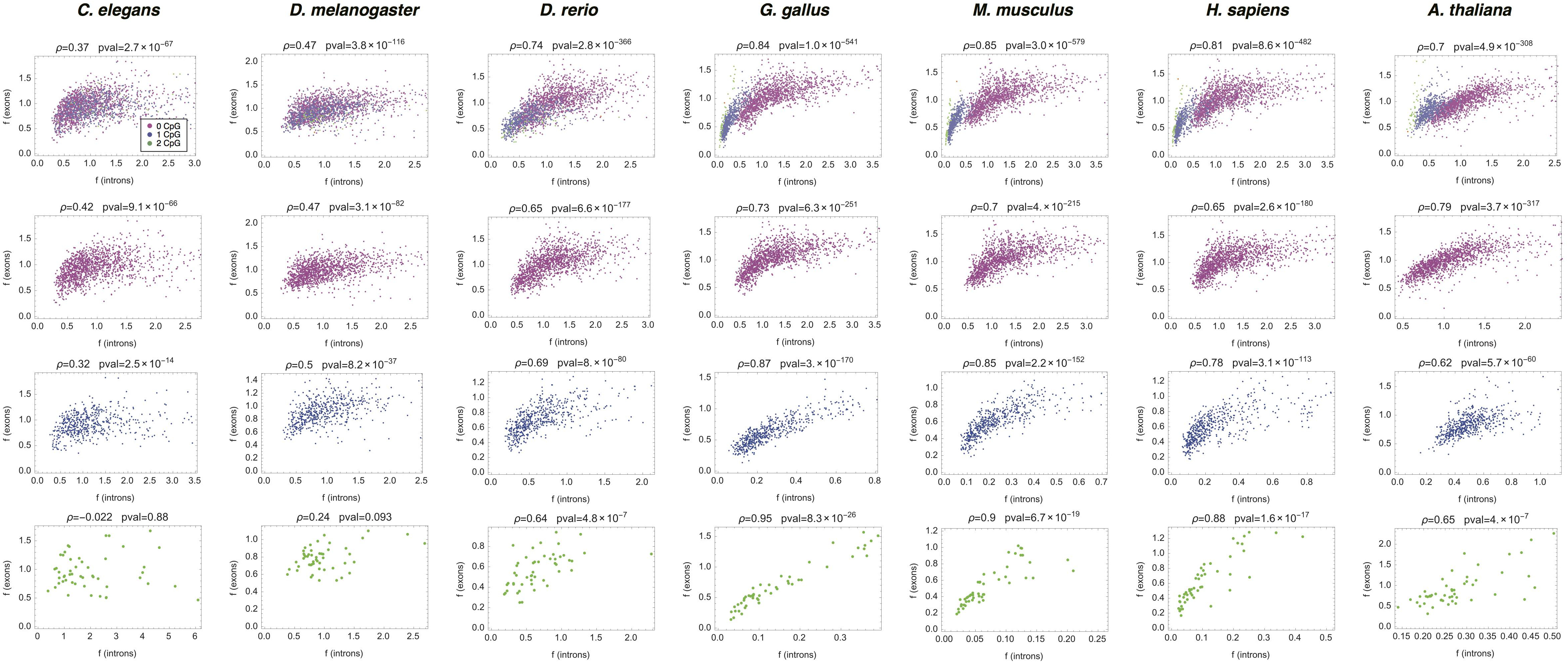
Correlation of word frequencies *f*_*i*_ measured in exons vs. in introns for different species. Top row shows results for all 6-mers, colored by CpG content (see legend); bottoms three rows plot separately within each CpG category. Spearman correlation ρ and p-value are indicated in each plot. Word frequencies *f*_*i*_ were computed relative to 1-mer null (introns) and relative to synonymous codon shuffling statistics (exons).

**Figure S9:**
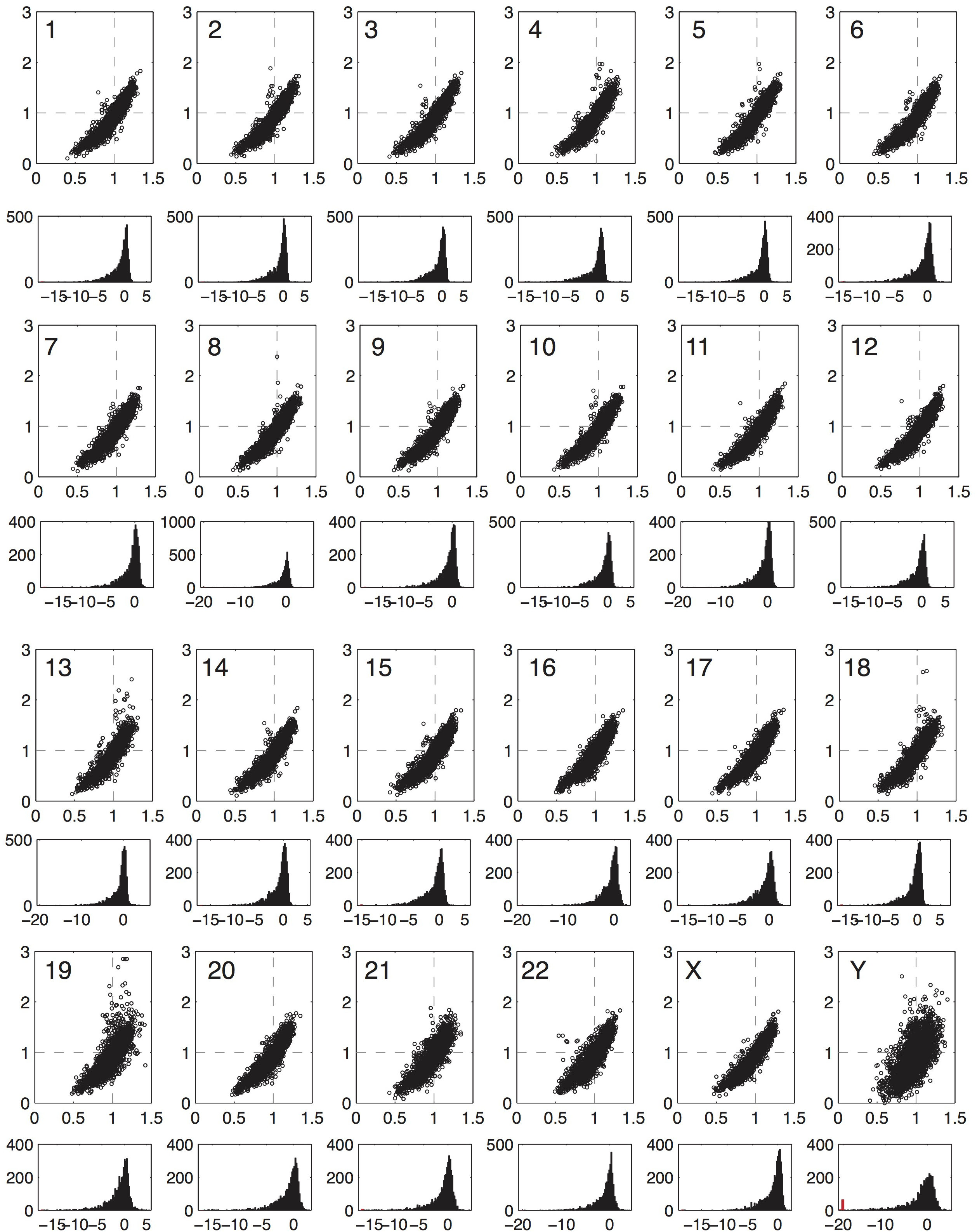
Word frequency plots of exons for all human chromosomes. For each chromosome, the top panel shows the 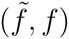 plot, and the bottom panel gives the excess pressure distribution, with ∆*s/u* on the *x*-axis. Codon shuffling was performed separately for each chromosome.

**Figure S10:**
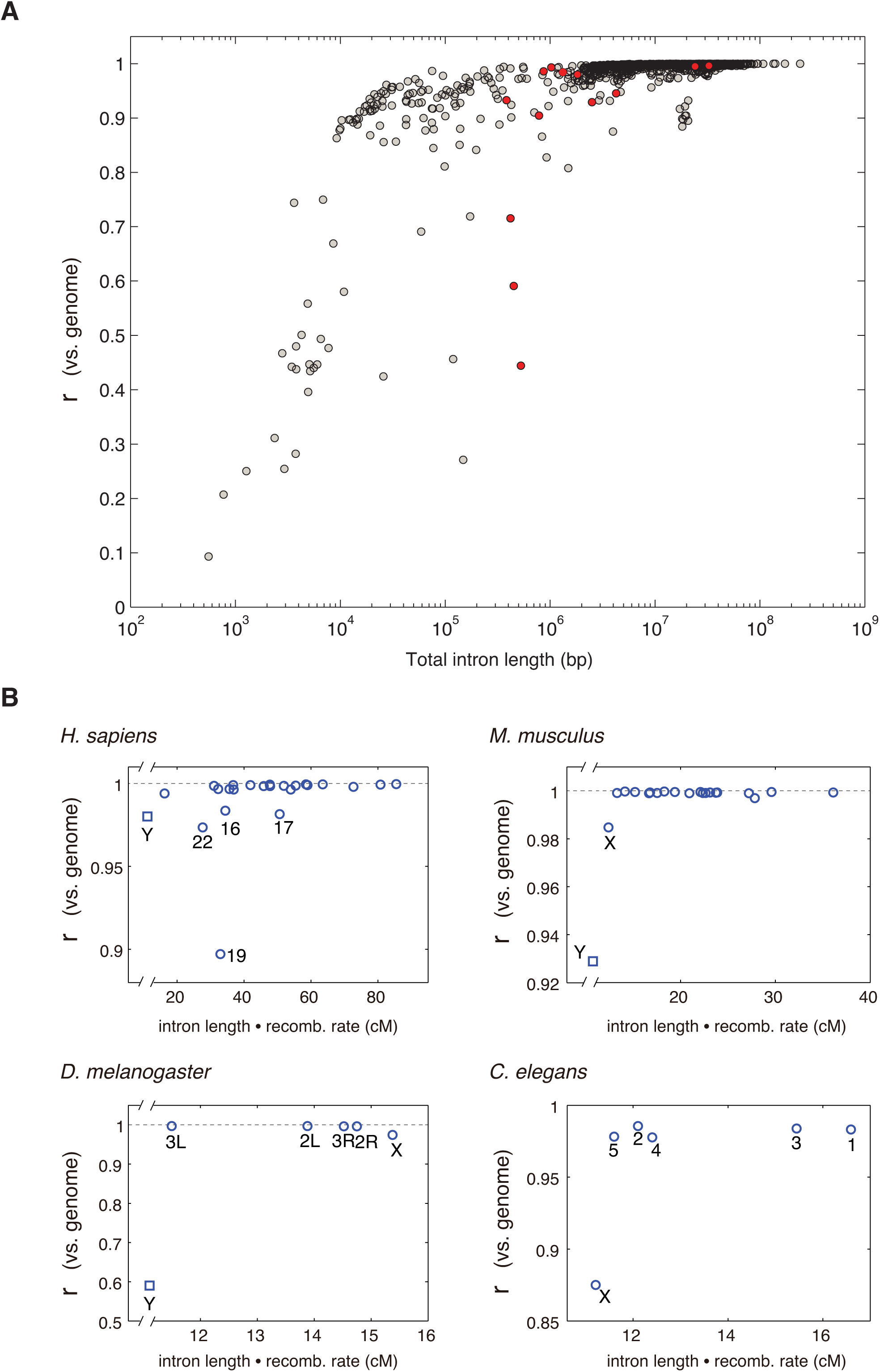
Relation of physical size and recombination rates to chromosomal word frequency deviations. **A**. Using intron data of each chromosome in the eukaryotic dataset (1,304 chromosomes), we compute the correlation coefficient of its relative word-frequency vector *f* vs. the genome-wide average, and plot this value against the chromosome’s total intron length. Relative word frequencies of each chromosome were calculated with respect to the 1-mer null, i.e. expected frequencies based on chromosome-specific nucleotide composition; the genome-wide average was relative to the genome-wide nucleotide composition. The plot indicates that the relative word composition of smaller chromosomes tends to correlate less strongly with the rest of the genome than that of larger chromosomes. Y / W chromosomes are indicated as red points. **B**. Subplots show model species in which chromosomal recombination rates have been measured, showing the correlation coefficient of each chromosome’s *f* vector vs. the genome-wide average. Relative word frequencies were computed as in panel (A). Y chromosomes are shown separately as squares. The x-axis was calculated as the recombination rate (cM / Mb) times the total length of introns analyzed (Mb). Recombination rates were obtained from [27] (human and mouse) and [28] (fly). In *C.elegans*, each chromosome crosses over exactly one time per meiosis, which corresponds to 50 cM [29]. We note that the further deviation of human chromosome 19 is potentially related to the prevalence of large, intrachromosomal duplications in its evolutionary history [30].

**Figure S11:**
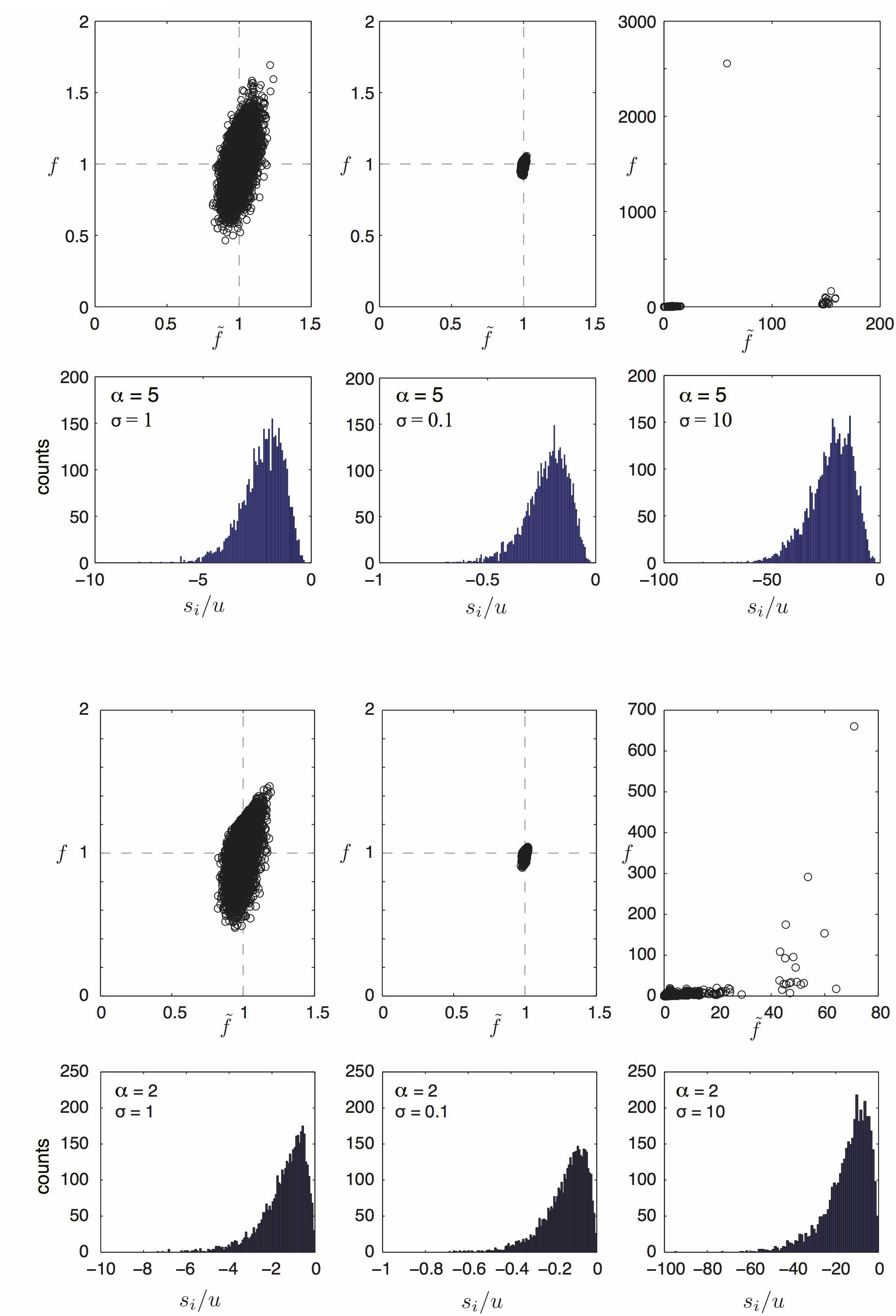
Model solution using Gamma-distributed random pressures, with given values of the shape parameter *α* and the standard deviation *σ*. The values of - *s*_*i*_ were assigned from the corresponding Gamma distribution, and used in the model (Eq. 2.2) to obtain the equilibrium word frequency vector f. Upper panels show the 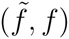 plots; lower panels show the histogram of selective coefficients in units of the mutation rate. All other details are as in Fig. 1.

**Figure S12:**
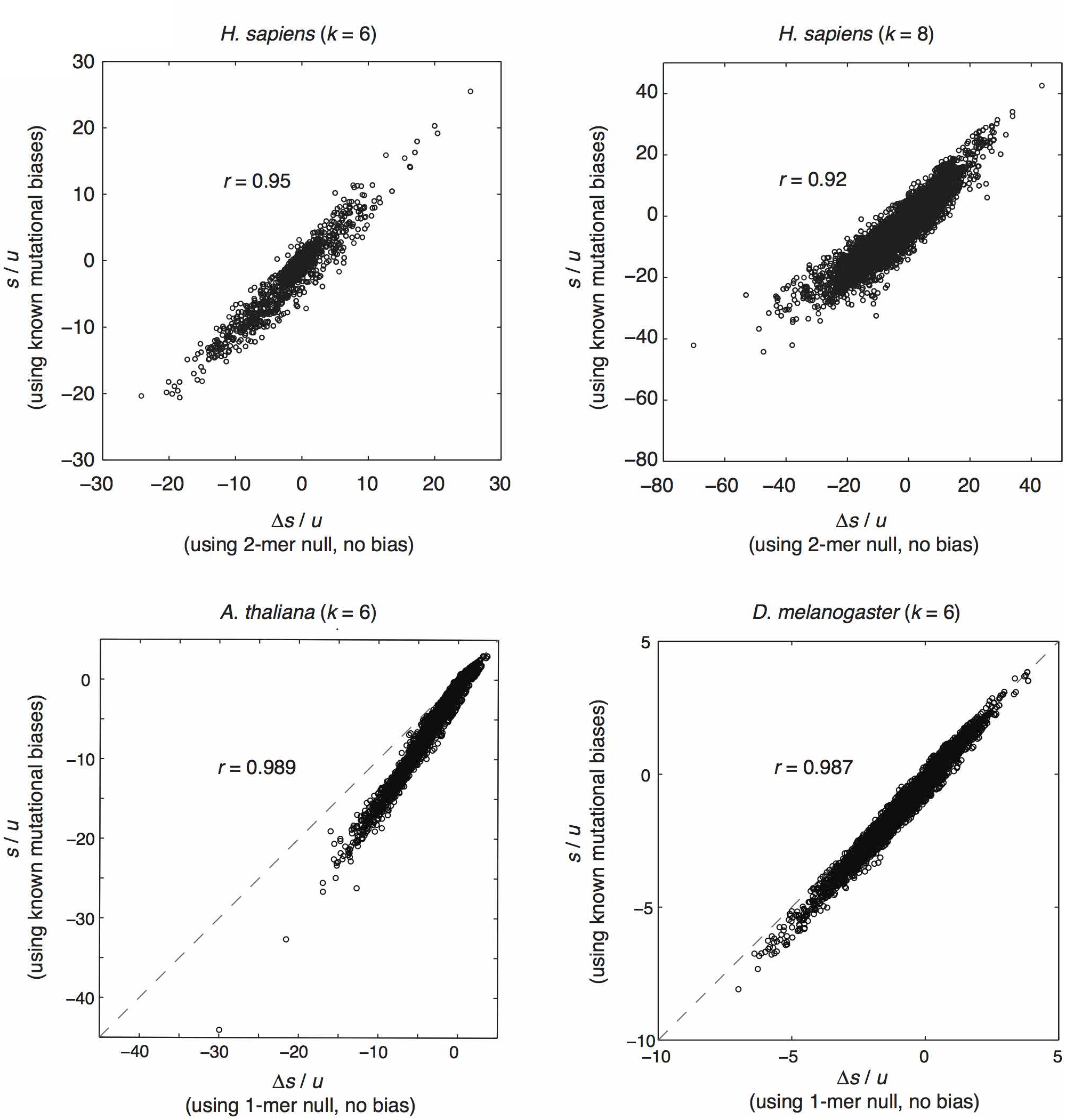
Comparison of selective coefficients inferred using known mutational biases vs. using measured nucleotide or dinucleotide composition for *H. sapiens, A. thaliana*, and *D. melanogaster*. To compute *s*_*i*_/*u* using known mutational biases, a mutation transition matrix *M*_*ij*_ was constructed using measured biases as follows. For the human genome, mutational biases measured in [15] were used, and the rate of dinucleotide CpG mutation was set to 11u as given in [17]. For plant and fly mutational biases, we used data from [13] and [12], respectively. Using the given mutational transition matrix in Eq. (2.5) and substituting in Eq. (2.4), we computed 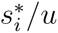 using Eq. (2.4) from the raw (unnormalized) frequencies *f*_*i*_ measured in introns. To compute excess pressures *∆s*_*i*_/*u*, we used an unbiased transition matrix and the unnormalized frequencies *f*_*i*_ to obtain *s*_*i*_, and used the expected frequencies 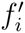 using the 2-mer null model (human) or 1-mer null model (plant and fly) to obtain 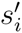, which yielded 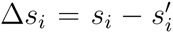. In all three cases, the plots show a very high correlation between the two different methods, which indicates that selective coefficients can be accurately inferred even in cases when mutational biases are not known, by using excess pressures relative to dinucleotide composition.

**Figure S13:**
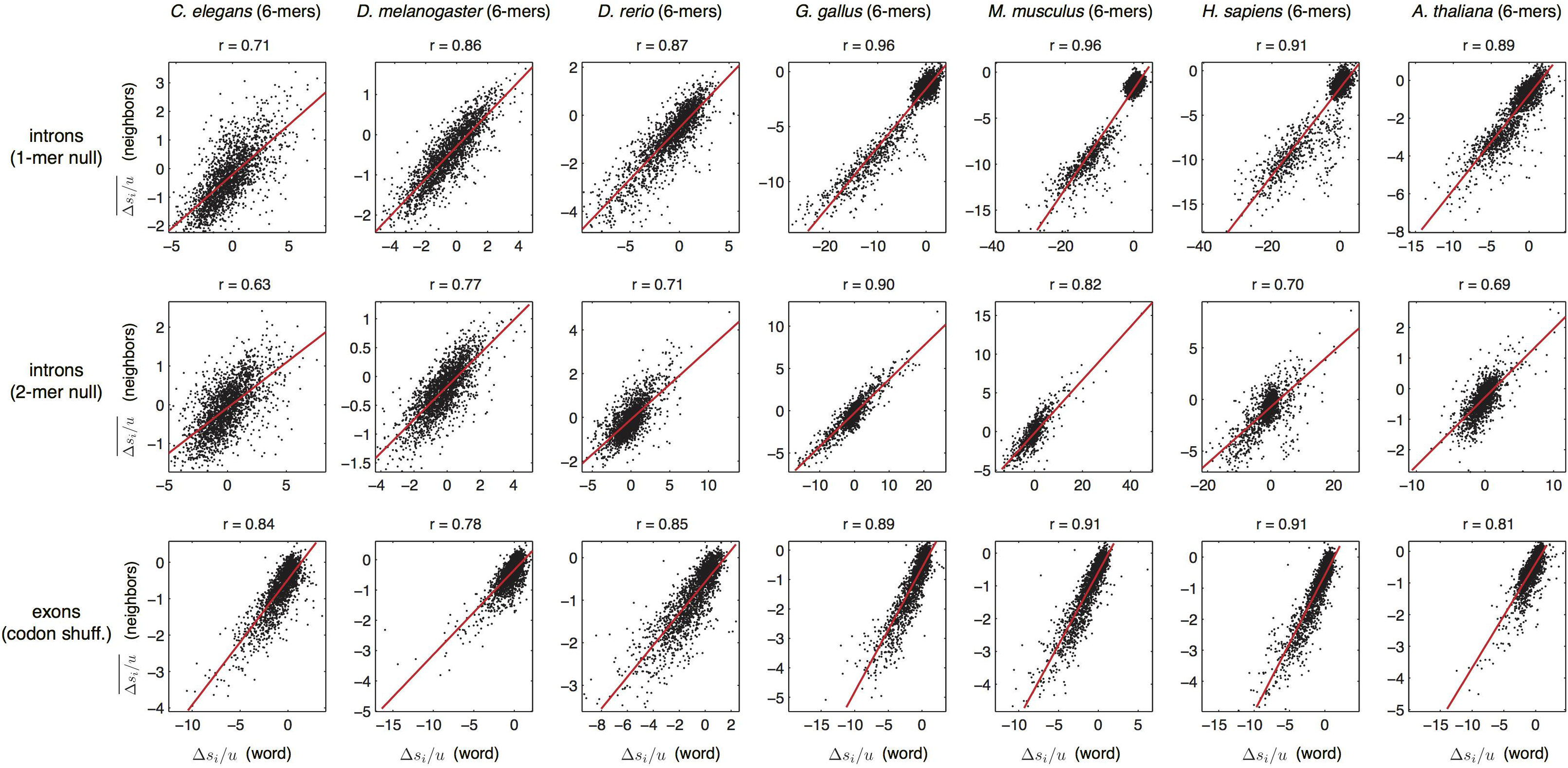
Correlation of excess pressures of words and their mutational neighbors in different species *(k* = 6). For each word, the excess pressure is plotted (*x*-axis) against the average excess pressure of its neighbors (*y*-axis). Excess pressures were calculated with respect to each of the null models (shown in separate rows). The correlation coefficient (*r*) from linear regression are given. All correlations are highly significant (p-val ≪ 10^−10^).

**Figure S14:**
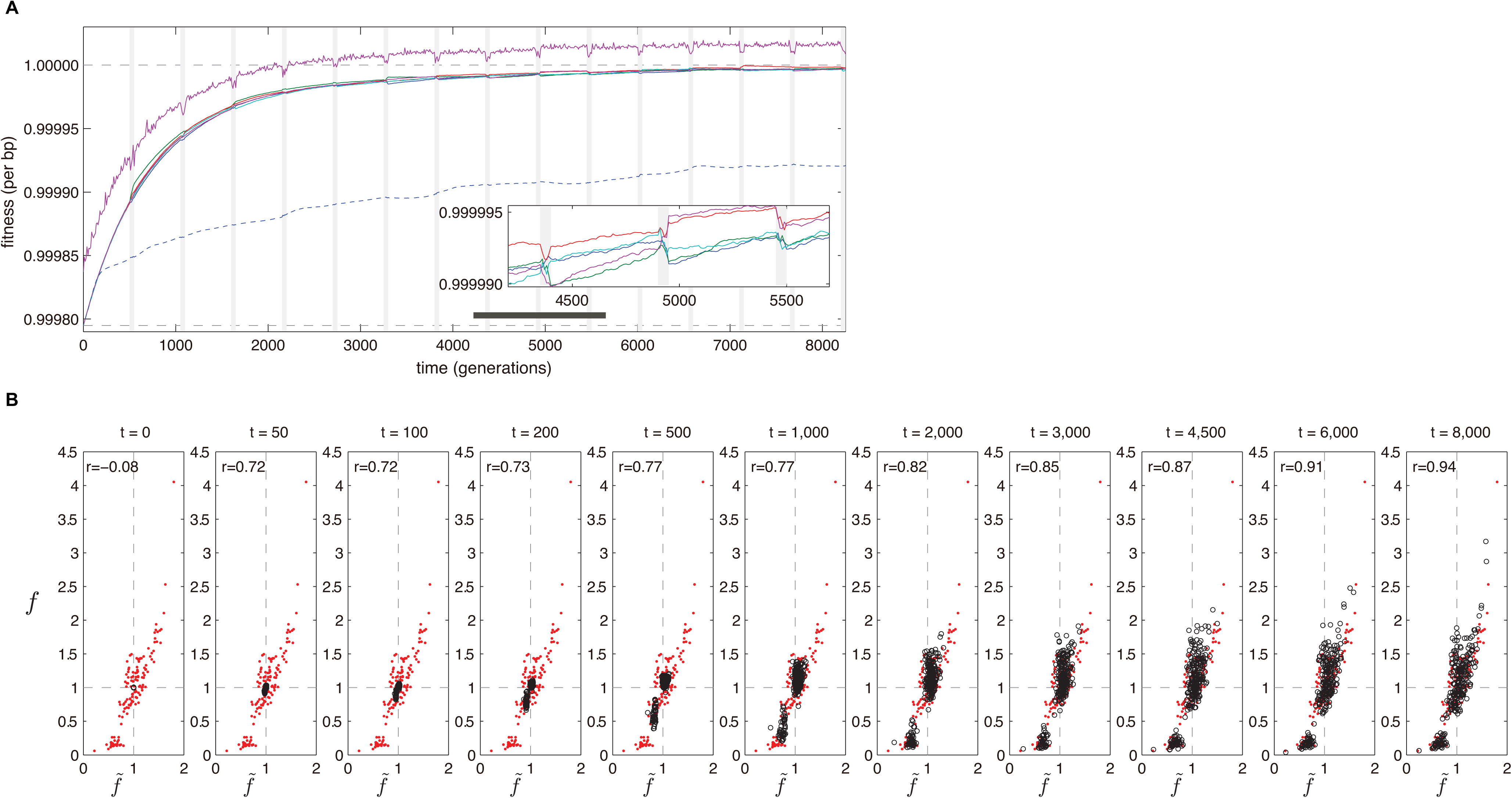
Dynamics of global per site fitness in simulated evolution with population bottlenecks (vertical gray bars) at regular intervals (see *Supplementary Methods*). **A**. Curves show the population average in five independent simulation runs with recombination (solid lines in color); the fittest sequence from one simulation with recombination (purple, top curve); and a run without recombination (dashed blue line). Dashed horizontal lines indicate the fitness for a uniform word composition (bottom line) and for the expected composition at mutation-selection balance (top line). Parameters: population size *N* = 10^5^ (10^3^ for bottlenecks); *L* = 2000bp; *u* = 1 x 10^−4^ per bp per generation; *s*_*i*_ are the measured values of *s*_*i*_/*u* from human introns using *k* = 4, multiplied by 10^−4^. Bottlenecks occur every 500 gen. and last 50 gen. Random pairs of sequences recombine with a cross-over length of 200bp. B. Snapshots of the evolution of the word frequency vector in the 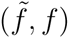 plane. Red dots correspond to the (4-mer) word frequencies in the human genome (i.e. the solution at mutation-selection balance) and the circles correspond to word frequencies in the simulated population at various time points.

**Figure S15:**
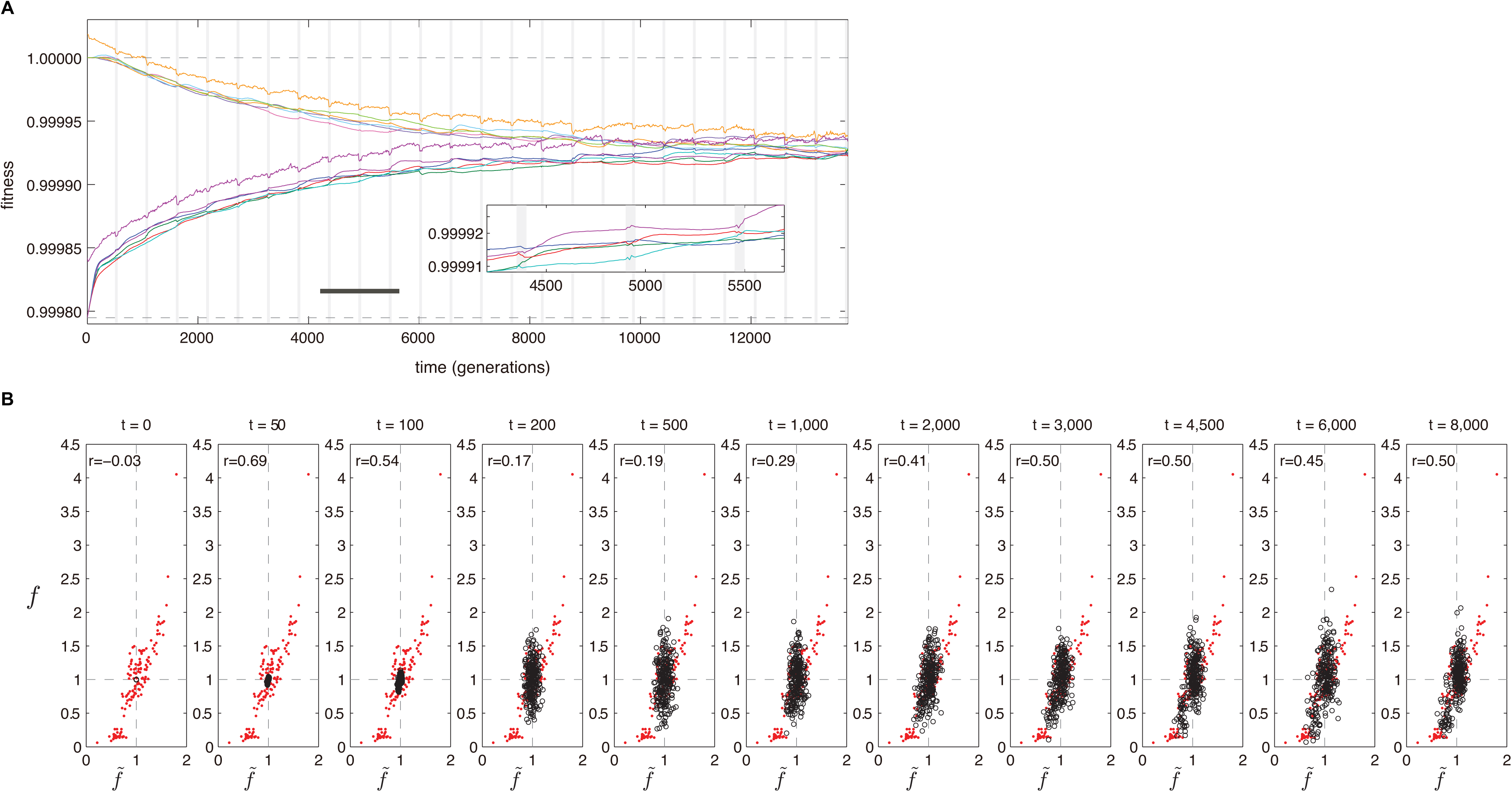
*In silico* evolution using human intron 4-mer pressures, showing the Muller’s ratchet effect in non-recombining populations. All parameters were the same as in Fig. S14, except that there was no recombination. One set of curves (lower) was initialized from random initial conditions; another set (upper) was initialized from the predicted equilibrium 4-mers frequencies. For each set, the top curve (purple or orange) corresponds to the maximally fit individual in the color-matched simulation run.

**Figure S16:**
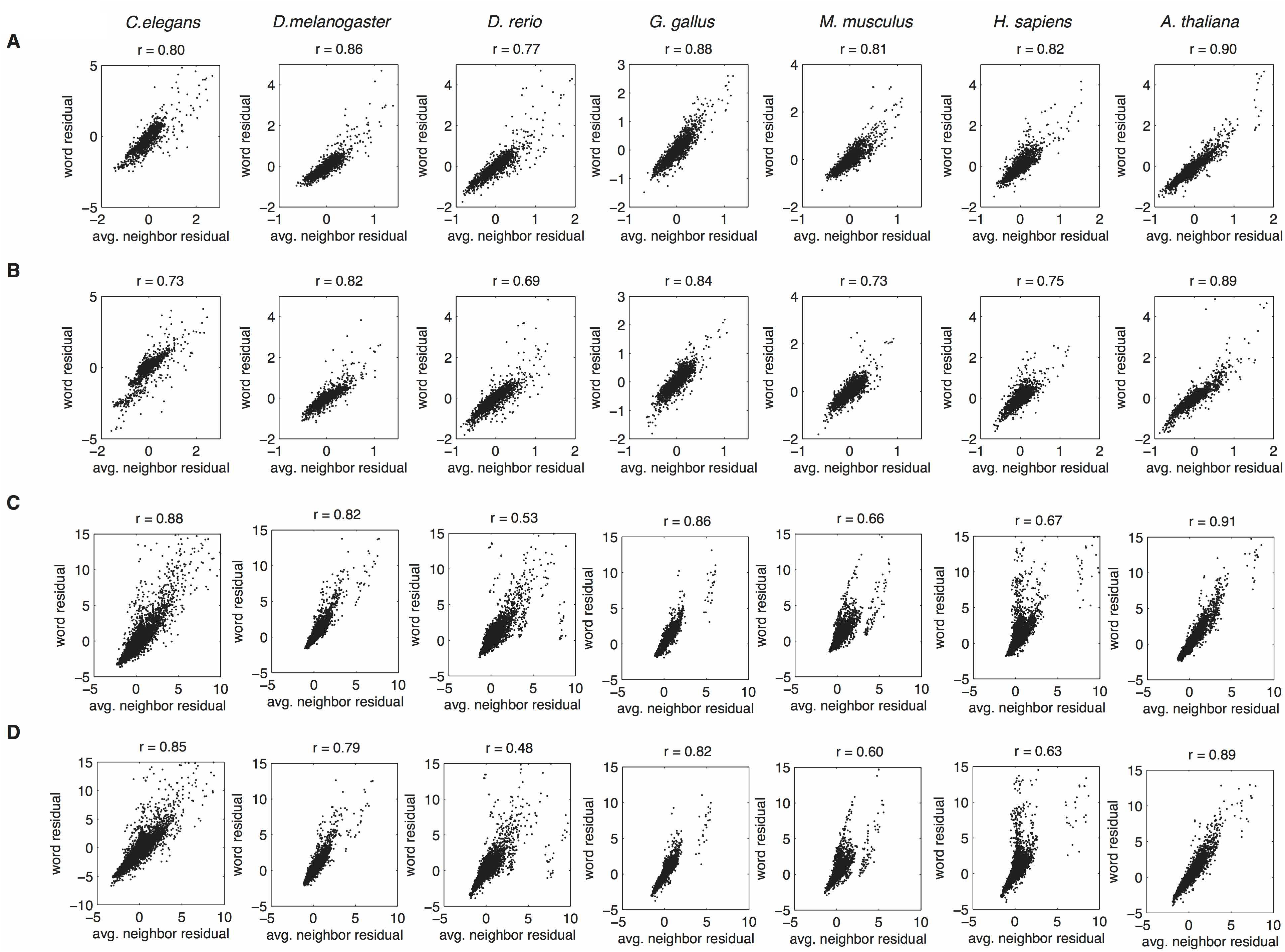
Word-neighbor plot of the residuals obtained from regression of word frequencies vs. word dinucleotide or trinucleotide composition (see Table S5 caption). Panels correspond to regression of 6-mer frequencies on dinucleotide composition (A) and trinucleotide composition (B); and 8-mer frequencies on dinucleotide composition (C) and trinucleotide composition (D). For each word, we plot its residual on the vertical axis vs. the average residuals of its mutational neighbors on the horizontal axis. Pearson correlation *r* is given for each plot. The figure demonstrates that even when dinucleotide or trinucleotide biases have been regressed out of the word frequency data, strong word-neighbor correlations persist.

**Figure S17:**
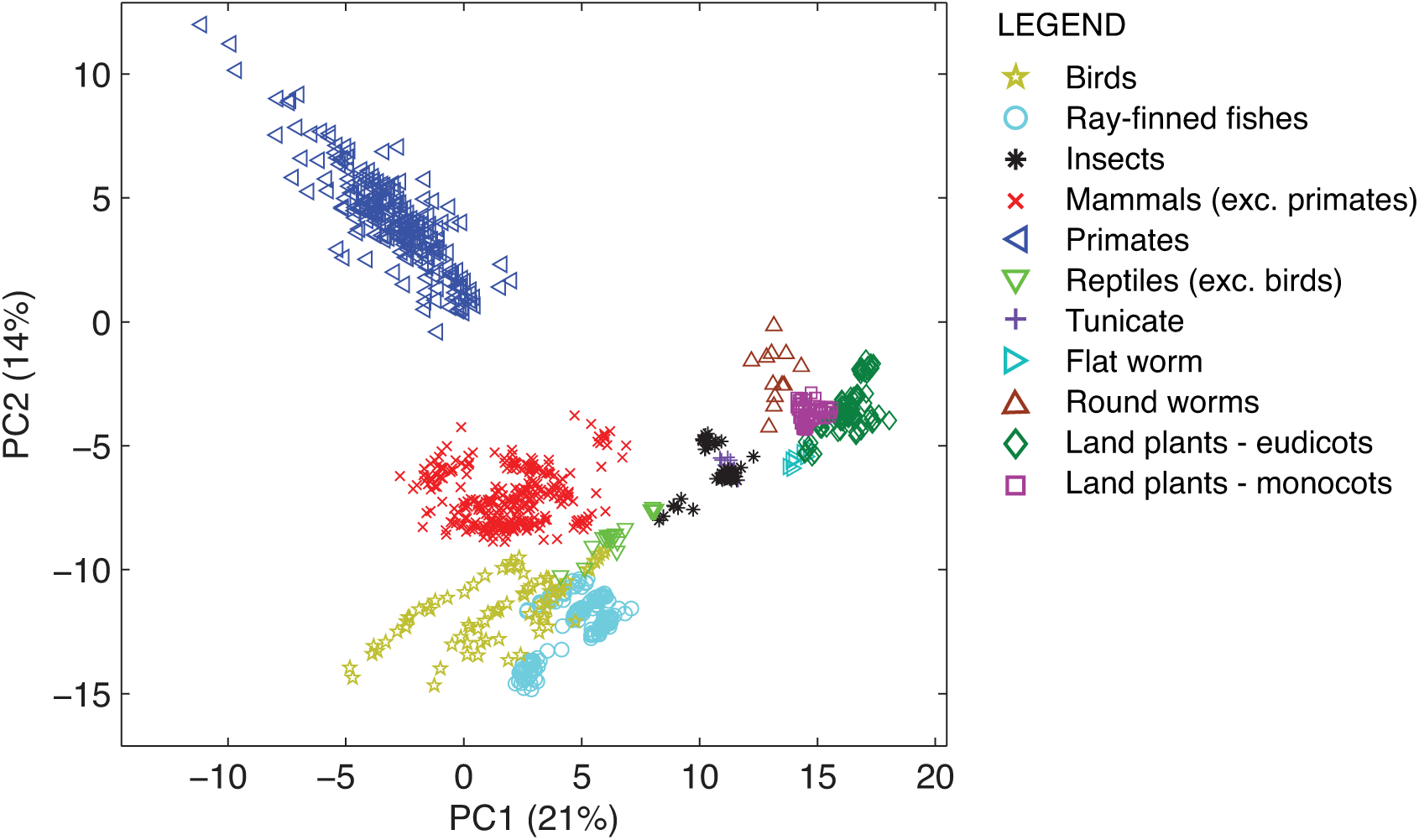
Word frequency PCA plot of all eukaryotic chromosomes from all phylogenetic groups, indicated by shapes & colors according to the legend. See Table S3 for further species information. The analysis was based on intronic 6-mer frequencies. Frequencies were normalized using the 2-mer null model.

**Figure S18:**
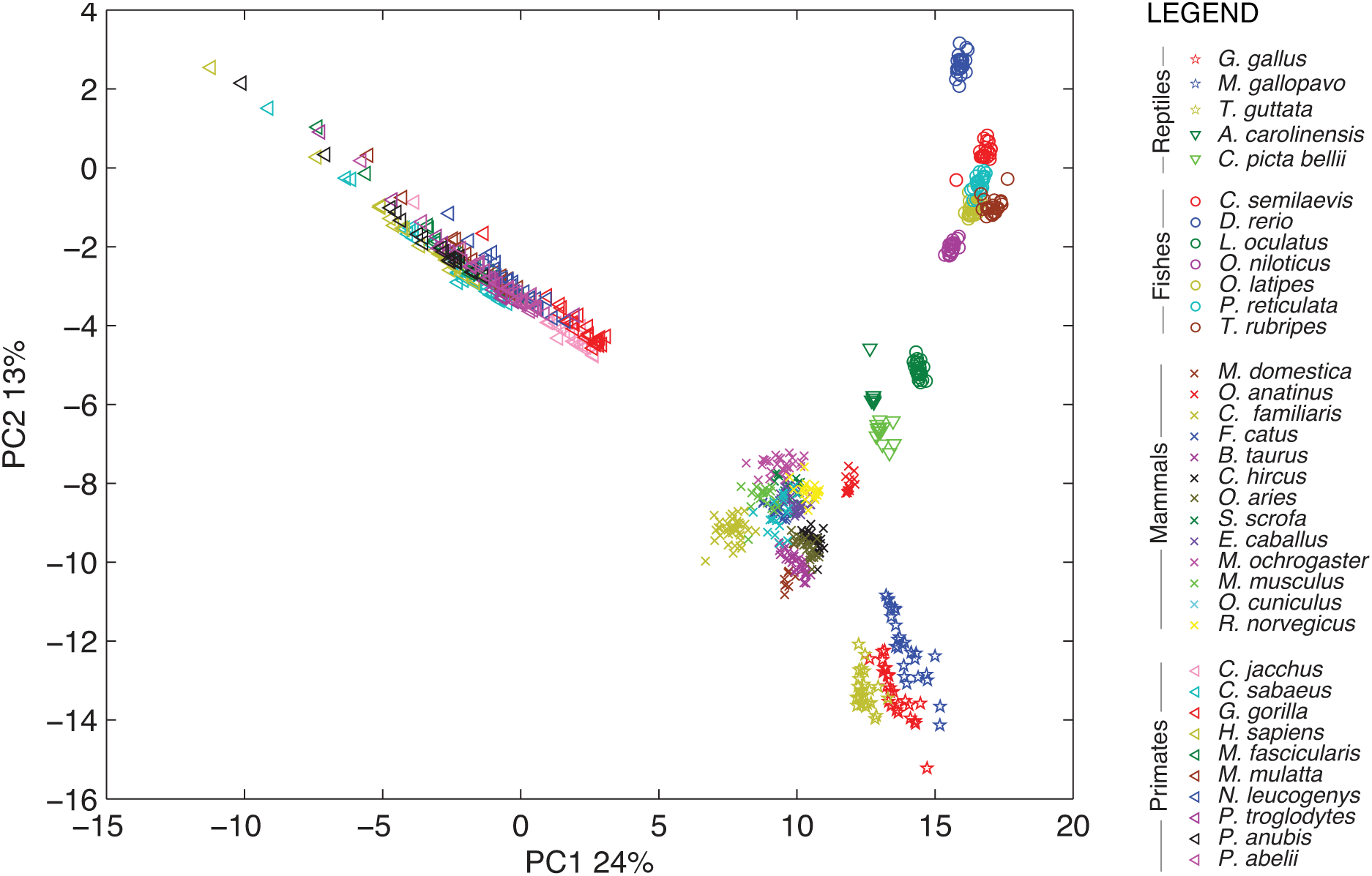
Word frequency PCA plot for all vertebrate chromosomes, showing the distinct separation of primates (left pointing triangles) away from the other mammals (x marks). See Table S3 for further species information. The analysis was based on intronic 6-mer frequencies. Frequencies were normalized using the 2-mer null model.

**Figure S19:**
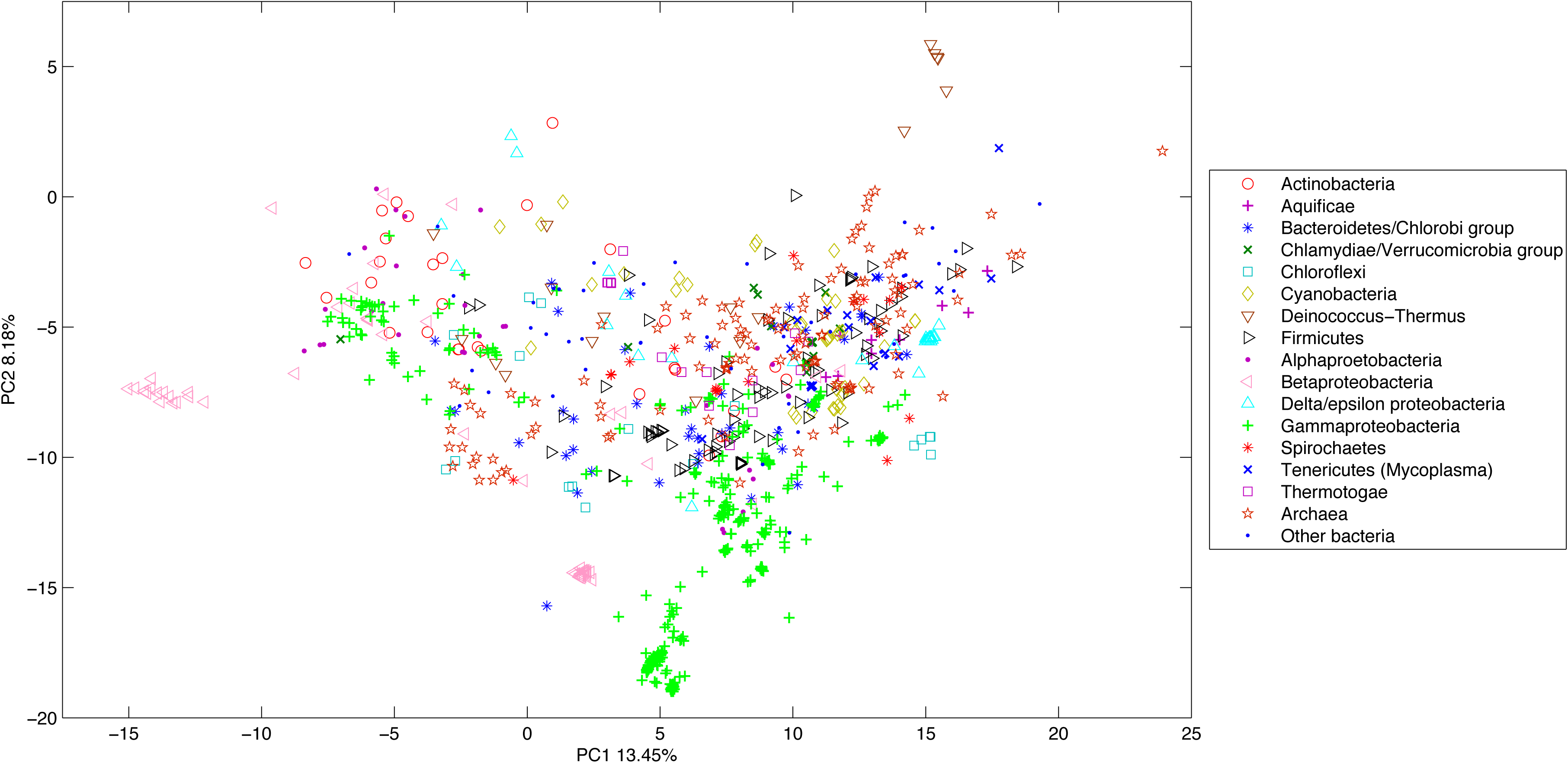
Word frequency PCA plot for all bacterial species (947 total). Color/symbol combinations denote different phyla (in the case of proteobacteria, further subdivisions are shown). Phyla with fewer than 20 genomes available were combined into a single group called ‘other bacteria’. The analysis was based on exonic 6-mer frequencies. Frequencies were normalized by synonymous codon shufflings.

**Figure S20:**
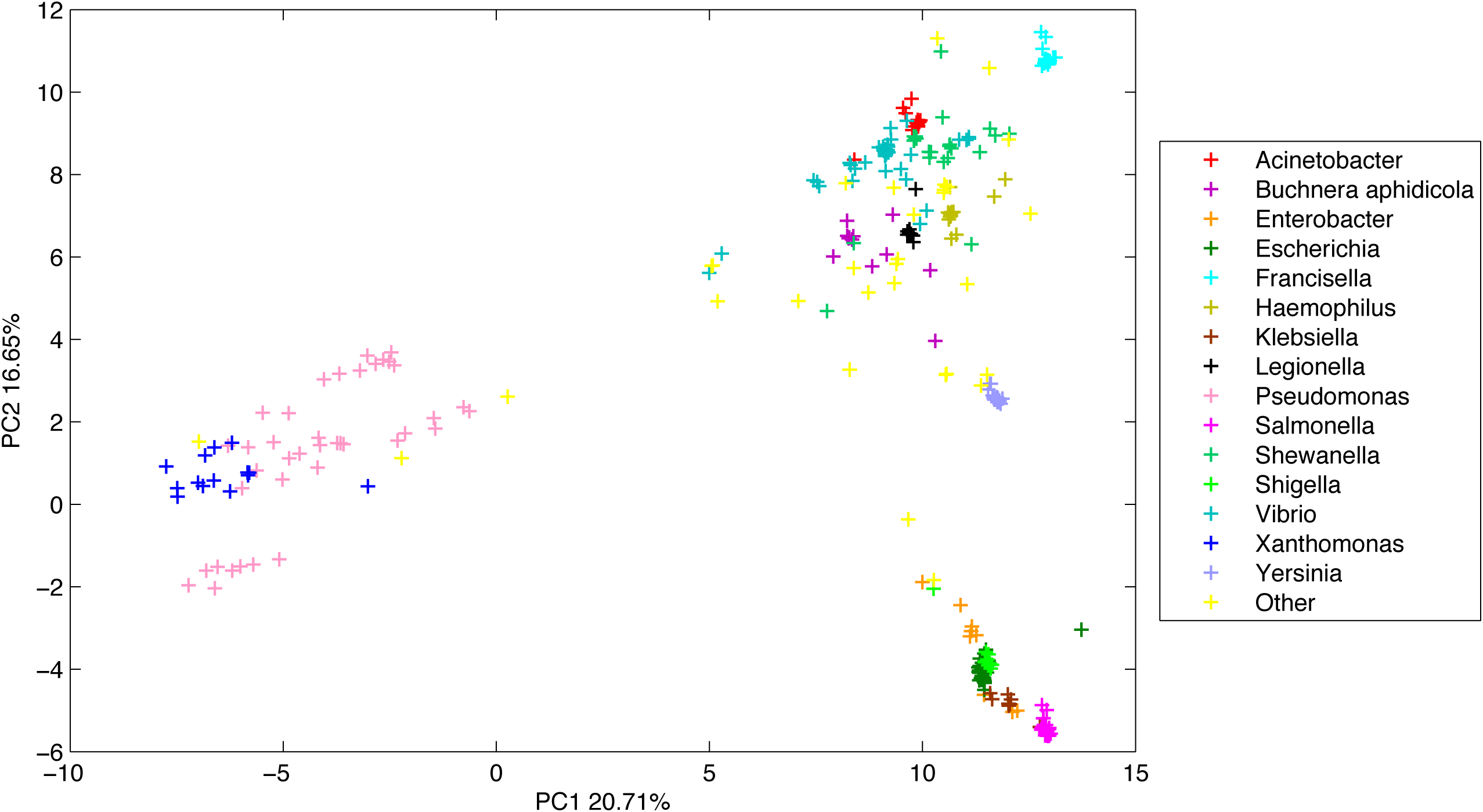
Word frequency PCA plot for β-Proteobacteria (356 species). Each genus is denoted by a different color. In several genus (Acinetobacter, Buchnera, Salmonella, Escherichia, Legiononella) clusters appear extremely tight because they consist mostly of strains from a single species; however other clusters consisting of multiple species are likewise relatively tight. The analysis was based on exonic 6-mer frequencies. Frequencies were normalized by synonymous codon shufflings.

**Figure S21:**
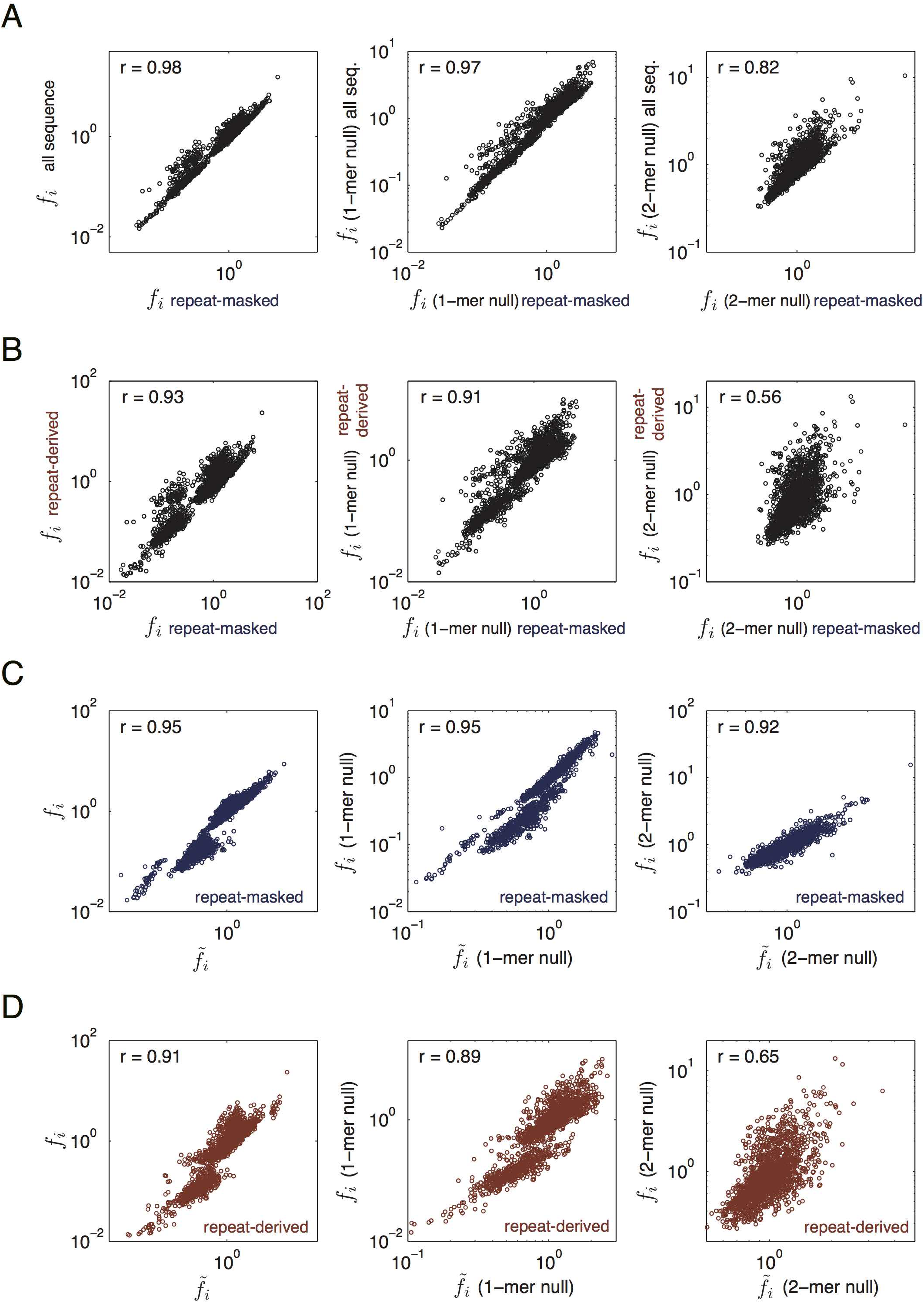
Word frequency distributions in the repeat and the non-repeat regions in introns of the human genome. Repeat sequences were extracted from intron regions from the repeat-masked sequence files (.mfa) in the RefSeq database, which uses the RepeatMasker program to identify repetitive or repeat-derived elements in the human genome. Word statistics were obtained separately from all repeat-derived or all repeat-masked segments of introns. The total length of repeat sequences extracted was 381.8Mb, leaving 454.2Mb of non-repeat intron sequences. (A, B) Correlations of word frequencies measured across all introns vs. across all repeat-masked (i.e. non-repeat-derived) intron regions (A), and across the repeat-derived intron regions vs. the repeat-masked regions (B). (C,D) Correlations between word and average neighbor frequencies measured in the repeat-masked regions (C) and in the repeat-derived regions (D). Plots from left to right are shown using raw word frequencies (normalized by the average word frequency), relative frequencies to the 1-mer null, and relative frequencies to the 2-mer null. To calculate relative frequencies, the 1-mer and 2-mer null models were constructed from nucleotide and di-nucleotide frequencies separately in the repeat-derived or repeat-masked portions accordingly. Pearson correlation coefficients (r) of log frequencies are indicated.

**Figure S22:**
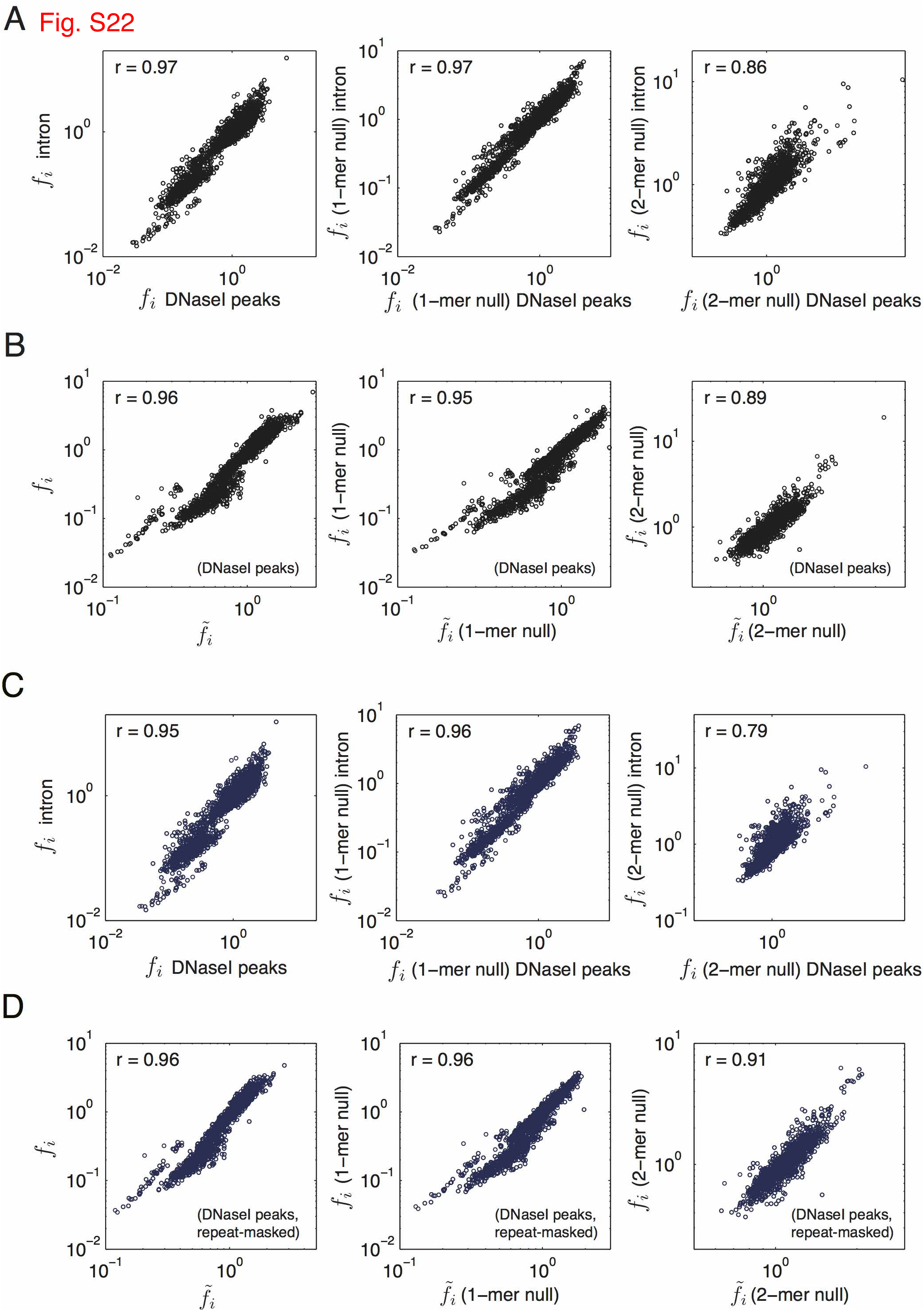
Word frequency distributions in DNase I hyper-sensitive regions in the human genome. DNase I hyper sensitive-regions (DNaseI peaks) were mapped by the UCSC ENCODE DNase analysis pipeline using the DNase-seq experimental data generated in [31]. DNaseI peaks were combined from 95 cell lines, and only the peaks in introns and intergenic regions were used (gene annotation was based on UCSC GENCODE v22). The total length of peak sequences analyzed was 431.4Mb. 6-mer frequencies and nucleotide and di-nucleotide frequencies were measured in each peak sequence and combined to calculate the genome-wide statistics. (A) Correlations of word frequencies obtained in DNaseI peaks vs. in the entire intronic regions. (B) Correlations between word and average neighbor frequencies in the DNaseI peaks. Pearson correlation coefficients (r) of log frequencies were indicated in the plots. (C, D) Same as panels (A, B) except that repeat-derived sequences (152.7Mb) were excluded from all the DNaseI peaks analyzed.

**Figure S23:**
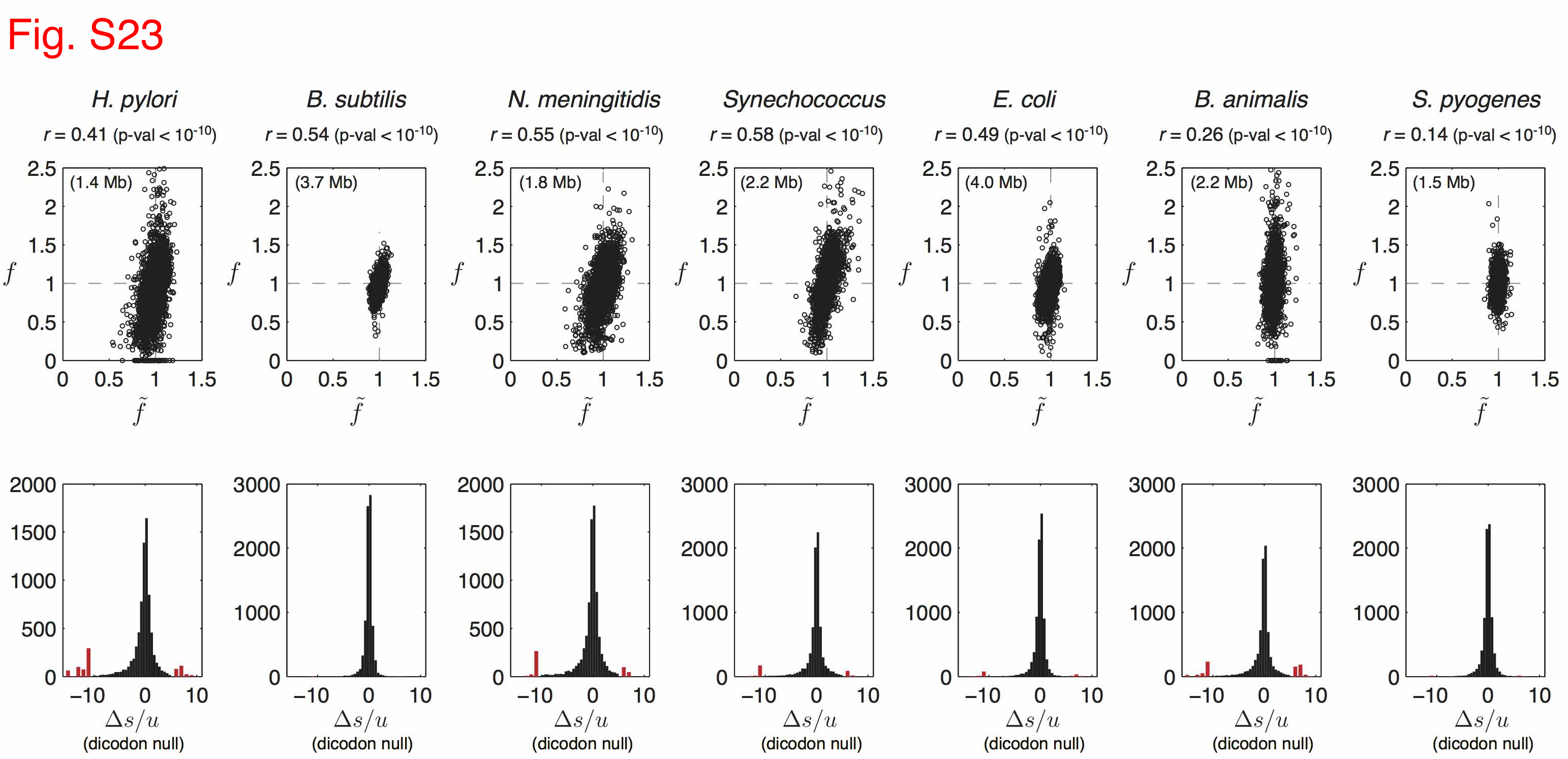
Word-neighbor plots for seven bacterial species relative to dicodon shuffling null model. For each species, the total exon sequence was used to construct 100 dicodon-shuffled randomizations, according to the shuffling scheme described in [32]. This scheme shuffles synonymous codons (i.e. maintaining amino-acid sequences) while preserving the dicodon pair statistics across the entire sequence. Since dicodons are 6 nucleotides long, this shuffling also preserves nucleotide, dinucleotide, trinucleotide, and tetranucleotide composition. Using the randomizations, we computed the expected word frequencies for all 7-mers. We used *k* = 7 instead of *k* = 6 to avoid any artifacts that could arise from the shuffling unit being of identical size to the *k*-mers we are measuring. Observed word frequencies in exon sequences are shown relative to the expected frequencies from the shuffled sequences (top row). Distributions of excess pressures relative to the dicodon shuffling null expectation are shown (bottom row). To show the extreme, red bins are used on the left and right of the distribution to indicate observations outside the range of (-10, 5): on the left, red bins correspond to excess pressures in the ranges <-1000, (-1000, −500), (-500,-100), (-100,-50), (-50,-10) and on the right to (5, 10), (10, 50), (50, 100), (100, 500), and > 500. The following genomes were used in the above analyses: *Helicobacter pylori 26695, Bacillus subtilis 168, Neisseria meningitidis MC58, Synechococcus sp. WH8102, Escherichia coli K-12 W3110, Bifidobacterium animalis AD011, Streptococcus pyogenes M1 GAS*.

### 3.2 Supplemental Tables

**Table S1:**
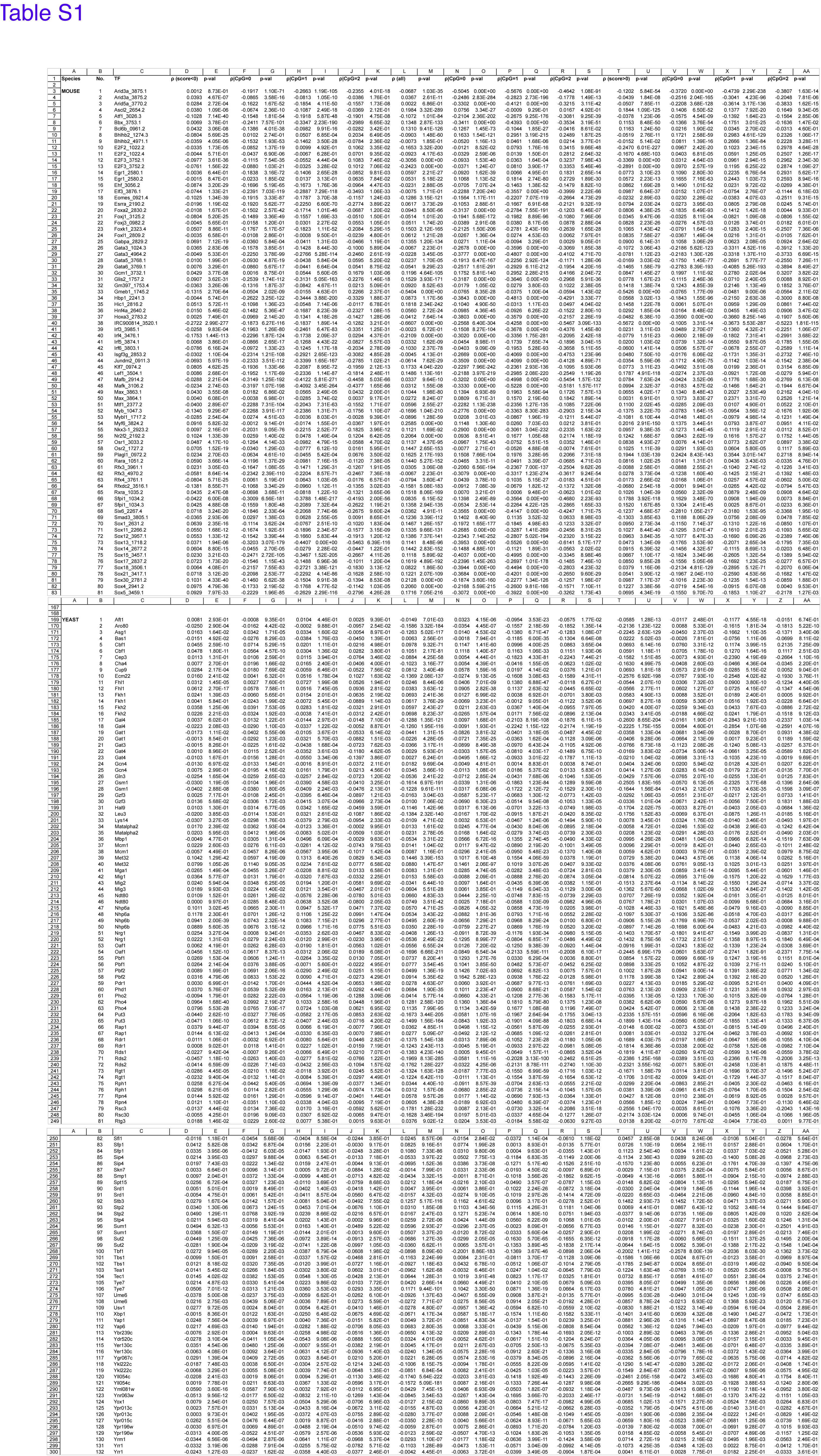
Transcription factor binding data correlated with genomic *k*-mer statistics in eukaryotic genomes. For each species and each TF, we computed the Spearman correlation (ρ) of binding scores *b*_*i*_ vs. genomic word frequencies *f*_*i*_, where *i* indexes over all 8-mer words. Frequencies *f*_*i*_ corresponded to the raw counts divided by expected counts based on nucleotide composition (i.e. the 1-mer null) for mouse, fly, worm, and human; counts were determined using all intron data in these four species. In yeast, which has few introns, the raw counts were made across all exons, and the expected counts were obtained from the synonymous codon shuffling scheme. Binding scores corresponded to z-scores for mouse [20] and fly datasets [21, 22, 23], and to e-values for worm [24] and yeast [25] datasets; the human dataset [26, 22] had 5 TFs with z-score data and 3 TFs with e-value data as indicated. We report ρ on three different subsets of *k*-mer words: (1) Columns D-K report ρ using all words *i* satisfying *b*_*i*_ < 0; (2) columns L-S use all words; and (3) columns T-AA use words satisfying *b*_*i*_ > 0. The p-value associated with each measurement of ρ is reported in the column immediately to its right. For each of the subsets 1-3, we report four measurements of ρ using (a) all words in the subset, (b) conditional on having 0 CpG, (c) 1 CpG, and (d) 2 CpG’s.

**Table S2:**
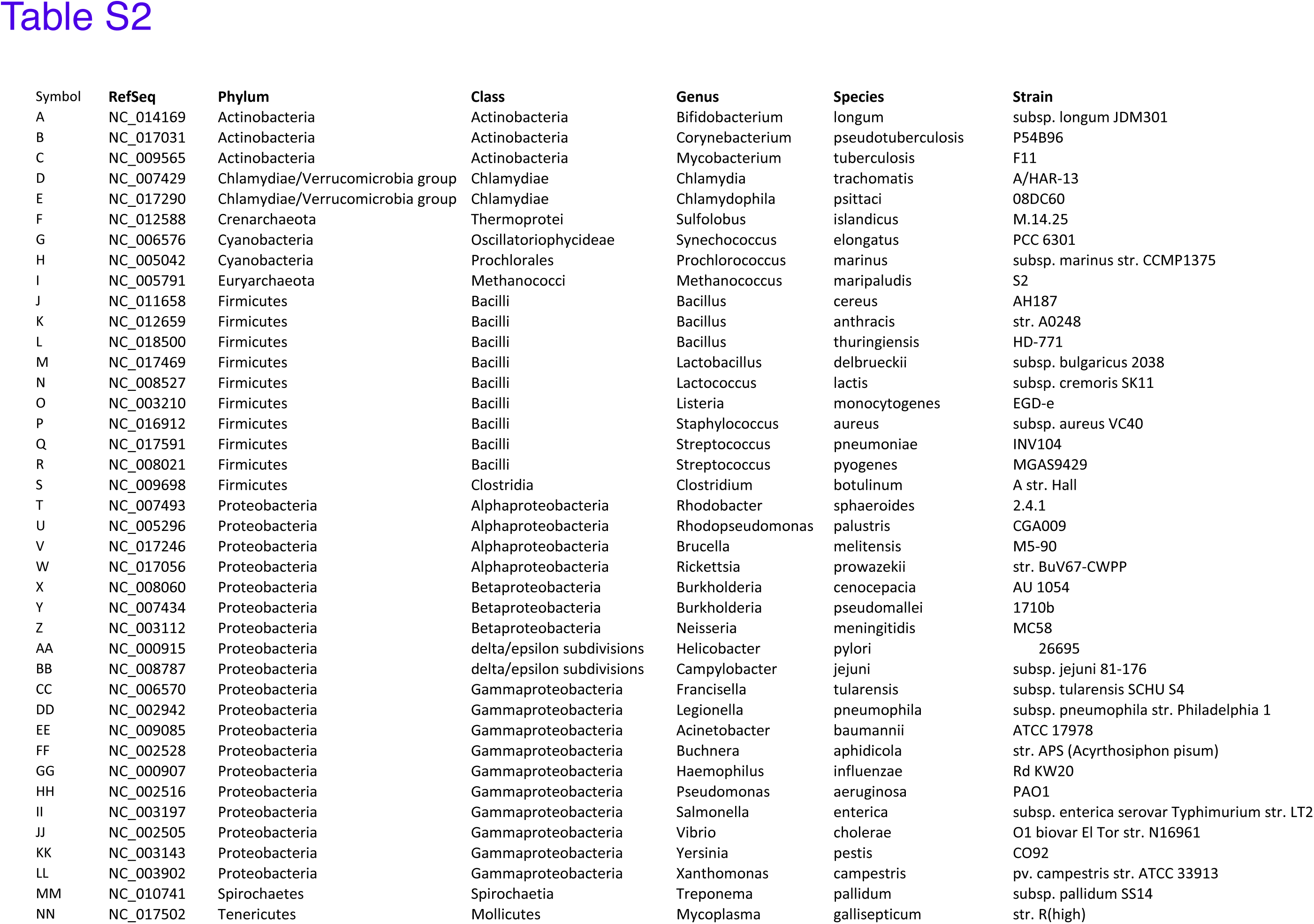
List of bacterial genomes and species analyzed.

**Table S3:**
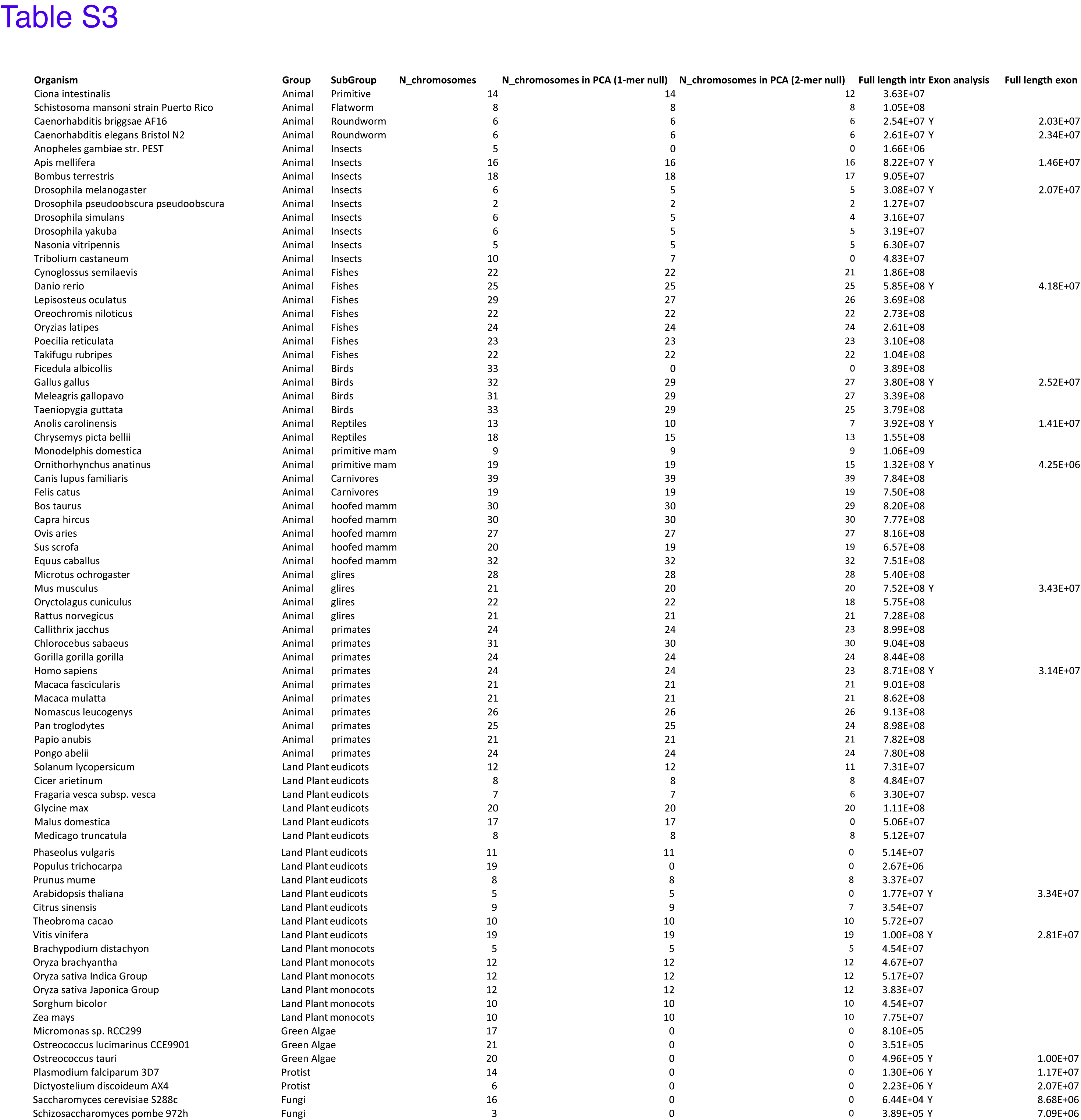
List of eukaryotic genomes and species analyzed.

**Table S4:**
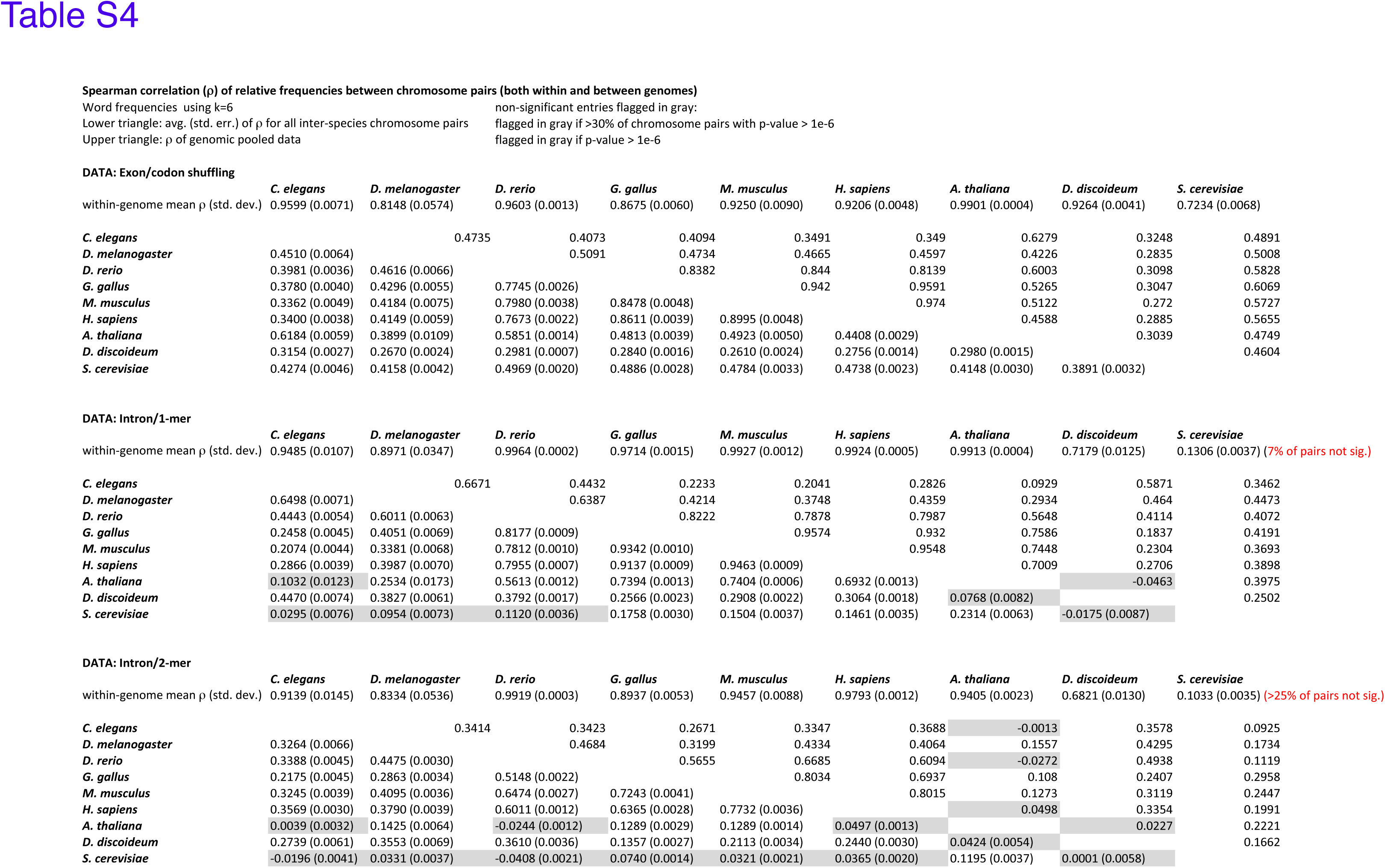
Spearman correlation coefficients (ρ) of relative word frequency vectors between chromosome pairs either within genomes or between eukaryotic species. Relative frequencies of 6-mers were measured in exons (using the codon shuffling scheme) and in introns (using the 1-mer and 2-mer null models). Within-species ρ values were computed for all chromosome pairs of the same species; average and standard deviations of ρ across chromosome pairs are reported. Between-species ρ values are shown in three matrices (corresponding to the three null models). For each pair of species, the upper triangle of each matrix indicates the correlation between genome-wide relative word frequencies, and the lower triangle gives the mean and standard error of ρ across all pairs of chromosomes.

**Table S5:** Analysis of variance (ANOVA) of relative word frequencies regressed on dinucleotide and trinucleotide composition in different genomes (see Secs. 1.4 and 2.2.1 for details).

| Species | $f$ | | $f$ (1-mer) | | $f$ (2-mer) | |
| --- | --- | --- | --- | --- | --- | --- |
| | $r^2$ | $r^2(\text{CpG})$ | $r^2$ | $r^2(\text{CpG})$ | $r^2$ | $r^2(\text{CpG})$ |
| *C.elegans* | 0.43 | 0.20 | 0.43 | 0.18 | 0.13 | 0.043 |
| *D.melanogaster* | 0.63 | 0.33 | 0.54 | 0.16 | 0.030 | 0.014 |
| *D.rerio* | 0.49 | 0.35 | 0.47 | 0.38 | 0.13 | 0.11 |
| *G.gallus* | 0.64 | 0.47 | 0.67 | 0.57 | 0.11 | 0.083 |
| *M.musculus* | 0.58 | 0.48 | 0.63 | 0.58 | 0.047 | 0.036 |
| *H.sapiens* | 0.54 | 0.39 | 0.58 | 0.52 | 0.074 | 0.057 |
| *A.thaliana* | 0.67 | 0.47 | 0.64 | 0.43 | 0.075 | 0.049 |

6-mer data regressed on 3-mer counts
| Species | $r^2$ | | | |
| --- | --- | --- | --- | --- |
| | $f$ | $f$ (1-mer) | $f$ (2-mer) | $f$ (3-mer) |
| *C.elegans* | 0.60 | 0.56 | 0.41 | 0.079 |
| *D.melanogaster* | 0.73 | 0.66 | 0.42 | 0.014 |
| *D.rerio* | 0.62 | 0.62 | 0.45 | 0.066 |
| *G.gallus* | 0.75 | 0.77 | 0.42 | 0.17 |
| *M.musculus* | 0.70 | 0.70 | 0.16 | 0.087 |
| *H.sapiens* | 0.67 | 0.66 | 0.32 | 0.067 |
| *A.thaliana* | 0.74 | 0.72 | 0.39 | 0.16 |

8-mer data regressed on 2-mer counts
| Species | $f$ | | $f$ (1-mer) | | $f$ (2-mer) | |
| --- | --- | --- | --- | --- | --- | --- |
| | $r^2$ | $r^2(\text{CpG})$ | $r^2$ | $r^2(\text{CpG})$ | $r^2$ | $r^2(\text{CpG})$ |
| *C.elegans* | 0.30 | 0.14 | 0.15 | 0.075 | 0.054 | 0.024 |
| *D.melanogaster* | 0.43 | 0.23 | 0.34 | 0.11 | 0.030 | 0.017 |
| *D.rerio* | 0.19 | 0.15 | 0.18 | 0.15 | 0.077 | 0.067 |
| *G.gallus* | 0.49 | 0.36 | 0.55 | 0.47 | 0.095 | 0.082 |
| *M.musculus* | 0.32 | 0.26 | 0.34 | 0.32 | 0.0044 | 0.0034 |
| *H.sapiens* | 0.29 | 0.21 | 0.29 | 0.26 | 0.030 | 0.025 |
| *A.thaliana* | 0.52 | 0.38 | 0.50 | 0.32 | 0.046 | 0.032 |

8-mer data regressed on 3-mer counts
| Species | $r^2$ | | | |
| --- | --- | --- | --- | --- |
| | $f$ | $f$ (1-mer) | $f$ (2-mer) | $f$ (3-mer) |
| *C.elegans* | 0.43 | 0.20 | 0.14 | 0.033 |
| *D.melanogaster* | 0.50 | 0.42 | 0.21 | 0.016 |
| *D.rerio* | 0.25 | 0.25 | 0.17 | 0.050 |
| *G.gallus* | 0.58 | 0.65 | 0.19 | 0.12 |
| *M.musculus* | 0.39 | 0.39 | 0.012 | 0.011 |
| *H.sapiens* | 0.36 | 0.33 | 0.081 | 0.028 |
| *A.thaliana* | 0.57 | 0.58 | 0.20 | 0.11 |

### 3.3 Additional Files

#### File S1

TF binding score and word frequency correlation plots for all analyzed data. For each transcription factor, we output a series of plots including (a) its reported primary binding motif, (b) its histogram of binding scores over all 8-mers, (c) the scatter plot of binding scores *b*_*i*_ vs. average mutational neighbor scores 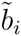, (d) a series of scatter plots showing *b*_*i*_ vs. *f*_*i*_ for words in different CpG categories (mouse, fly, human), or for all words (worm and yeast). Spearman correlation values ρ are shown, and significant values with *p* < 10^−6^ are indicated as ***. These plots were produced from the same data that was used to produce Table S1 (see caption for references).

## 4 Supplemental References

